# Integrative Multiscale Biochemical Mapping of the Brain via Deep-Learning-Enhanced High-Throughput Mass Spectrometry

**DOI:** 10.1101/2023.05.31.543144

**Authors:** Yuxuan Richard Xie, Daniel C. Castro, Stanislav S. Rubakhin, Timothy J. Trinklein, Jonathan V. Sweedler, Fan Lam

**Affiliations:** Department of Bioengineering, University of Illinois Urbana-Champaign, Urbana, IL 61801, United States; Beckman Institute for Advanced Science and Technology, University of Illinois Urbana-Champaign, Urbana, IL 61801, United States; Department of Molecular and Integrative Physiology, University of Illinois Urbana-Champaign, Urbana, IL 61801, United States; Department of Chemistry, University of Illinois Urbana-Champaign, Urbana, IL 61801, United States; Department of Electrical and Computer Engineering, University of Illinois Urbana-Champaign, Urbana, IL 61801, United States; Carle-Illinois College of Medicine, University of Illinois Urbana-Champaign, Urbana, IL 61801, United States; Carl R. Woese Institute for Genomic Biology. University of Illinois Urbana-Champaign, Urbana, IL 61801, United States

## Abstract

Elucidating the spatial-biochemical organization of the brain across different scales produces invaluable insight into the molecular intricacy of the brain. While mass spectrometry imaging (MSI) provides spatial localization of compounds, comprehensive chemical profiling at a brain-wide scale in three dimensions by MSI with single-cell resolution has not been achieved. We demonstrate complementary brain-wide and single-cell biochemical mapping via MEISTER, an integrative experimental and computational mass spectrometry framework. MEISTER integrates a deep-learning-based reconstruction that accelerates high-mass-resolving MS by 15-fold, multimodal registration creating 3D molecular distributions, and a data integration method fitting cell-specific mass spectra to 3D data sets. We imaged detailed lipid profiles in tissues with data sets containing millions of pixels, and in large single-cell populations acquired from the rat brain. We identified region-specific lipid contents, and cell-specific localizations of lipids depending on both cell subpopulations and anatomical origins of the cells. Our workflow establishes a blueprint for future developments of multiscale technologies for biochemical characterization of the brain.

## Introduction

Genomic and transcriptomic tools have transformed neuroscience by allowing us to visualize, untangle, and understand the spatiotemporal expression patterns of thousands of genes in the brain as well as how they are related to various functions and diseases^1–3^. Beyond gene expression profiles, the biochemical compositions and dynamics of metabolites, lipids, peptides and proteins play essential roles in many neurobiological processes^4,5^, and have been implicated in neurodevelopment^6^, learning, memory^7^, aging^8,9^, and a myriad of neurological or neurodegenerative diseases^10^. Approaches to characterize these molecular compositions offer invaluable insight complementary to transcriptomics. However, comprehensive biochemical profiling of both tissue and single cells at a whole-organ level remain challenging. Recent technical advances in single-cell measurements using isolated populations of individual cells and mass spectrometry (MS) have great potential to solve these bottlenecks, prompting single-cell metabolomics to be listed as one of the technologies to watch in 2023^11^. MS is recognized as a key method of choice for metabolomic and proteomic measurements due to its unique capability of untargeted, sensitive and specific detection of numerous biomolecules in both tissues^12,13^ and single cells^14–17^. Spatial organizations of biomolecules in the brain have been mapped at cellular and subcellular resolution using advanced mass spectrometry imaging (MSI) methods^18–21^. Nevertheless, small metabolites and lipid profiling of large brain regions in three dimensions at single-cell resolution with simultaneous brain-wide coverage and chemical detail (important for untargeted and unbiased molecular characterization) has yet been achieved. We provide several innovations to the existing workflows that enable multiscale biochemical profiling at a scale never attempted before. First, as existing high-resolution MSI is throughput limited, we integrate deep learning approaches to enhance the high-mass-resolving Fourier transform MS (FTMS) acquisition by 10-fold, enabling imaging of many tissue sections with brain-wide coverage and reconstruction of three-dimensional (3D) molecular distributions/atlases. Second, high-throughput single-cell MS (SCMS) allows populations of individual cells to be characterized^22^; however, isolated cells lack spatial context of tissue. We integrate both workflows (high-throughput tissue MSI and SCMS) to map the chemical profiles of single cells onto the tissue sections, allowing multiscale characterization of spatial-biochemical organization of the brain.

More specifically, we introduce MEISTER, a framework of mass spectrometry for integrative single-cell and tissue analysis with deep learning-based reconstruction that integrates high-throughput MS platforms with several technical innovations: (1) A deep-learning-based signal reconstruction approach capable of producing high-resolution mass spectra with significantly enhanced throughput for both tissue MSI and SCMS, (2) a multimodal image registration technique that produces coherent 3D reconstruction of MSI data from many tissue sections and affords quantitative analysis of regional chemical profiles, (3) a computational approach that exploits dictionary learning concepts to create and map cell-specific chemical profiles to tissue imaging data for multiscale integration. We validated MEISTER using computational simulations as well as experimental tissue MSI and SCMS data. With MEISTER, we achieved 3D mapping of the rat brain with an unprecedented combination of large volume coverage, high spatial resolution (50 𝜇m lateral and 16 𝜇m sections) over millions of pixels, and high chemical content (>1,000 lipid features). We also profiled 13,566 single cells that were isolated from five rat brain regions and built cell-type-specific chemical dictionaries, which were then mapped to the tissue images, to obtain spatially-resolved cell-type distributions across the brain. To further demonstrate the capabilities of our framework, we studied how lipids associate with the brain’s anatomical structures. We extracted lipid profiles from 11 brain regions by registering serial MSI sections to rat magnetic resonance imaging (MRI) brain atlas via a data-driven nonlinear image registration method that generated volumetric reconstruction of thousands of lipid features over a large brain volume while identifying region-specific lipid contents. With the single cell-to-tissue data integration approach, we identified heterogeneous lipid distributions and differential lipid features at both tissue and single-cell levels, discovering relationships of single-cell biochemical profiles to region-specific spatial distributions of lipids. We demonstrated the potential of MEISTER as a general multiscale tissue biochemical characterization approach by also applying it to another tissue type on rat pancreas and to molecules beyond lipids, e.g., peptides.

## Results

### MEISTER: A deep learning enabled, high-throughput multiscale MSI framework

MEISTER integrates high-throughput MS experiments, a deep-learning-based signal reconstruction method, and data-driven high-dimensional MSI analysis, to enable brain-wide, multiscale profiling of brain biochemistry. To resolve detailed chemical contents, we collected both high-resolution tissue MSI and SCMS data leveraging a high-throughput experimental platform using matrix-assisted laser desorption/ionization (MALDI) Fourier transform ion cyclotron resonance (FT-ICR) MS. Achieving brain-wide coverage and cell-specific profiling requires probing a large number of tissue sections and cells (Fig. 1a), which is time prohibitive on high-mass-resolution platforms such as FT-ICR (see Methods, signal modeling). To this end, we developed a deep-learning model to reconstruct high-resolution MS data from low-mass resolving measurements (Fig. 1b). In short, we model the underlying high-dimensional transient signals 𝑺 as points on a low-dimensional nonlinear manifold embedded in the high-dimensional space. These low-dimensional embeddings 𝒁 can be effectively learned by training a deep autoencoder (DAE) network, using experimental and/or simulated full transients with the desired mass resolution. The presence of low-dimensional representations implies that high-mass-resolution spectra can be reconstructed from significantly shorter transients than conventionally acquired. We realized this by training a “regressor” network jointly with the DAE to estimate the low-dimensional embeddings 𝒁 from only short transients 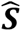, which were subsequently decoded (by the same decoder from the DAE) into high-mass-resolution data (More details in Extended Data Fig. 1a, and Methods). For 3D MSI, the networks were trained only on a small number of tissue sections and applied to reconstruct data for the remaining sections consisting of millions of pixels (Extended Data Fig. 1b, Methods). For SCMS, a small subset of cells were used for training and applied to large cell populations (Extended Data Fig. 1c), allowing a much higher data collection throughput. Particularly, an MSI data set containing more than 1.5 million pixels only required 20 h of acquisition time, which would have taken about 300 h using the conventional acquisition approach. This allowed us to image 16-µm-thick serial sections from rat brains that covers ∼10 mm range (along z) with a 50 µm raster width used in MSI. In conjunction with the MSI, 13,566 single cells isolated from five brain regions (the neocortex, hippocampus, thalamus, striatum, corpus callosum) were probed by an image-guided MALDI SCMS^23^ approach (Fig. 1a). Detailed experimental parameters for MSI and SCMS can be found in Methods.

**Figure 1.**
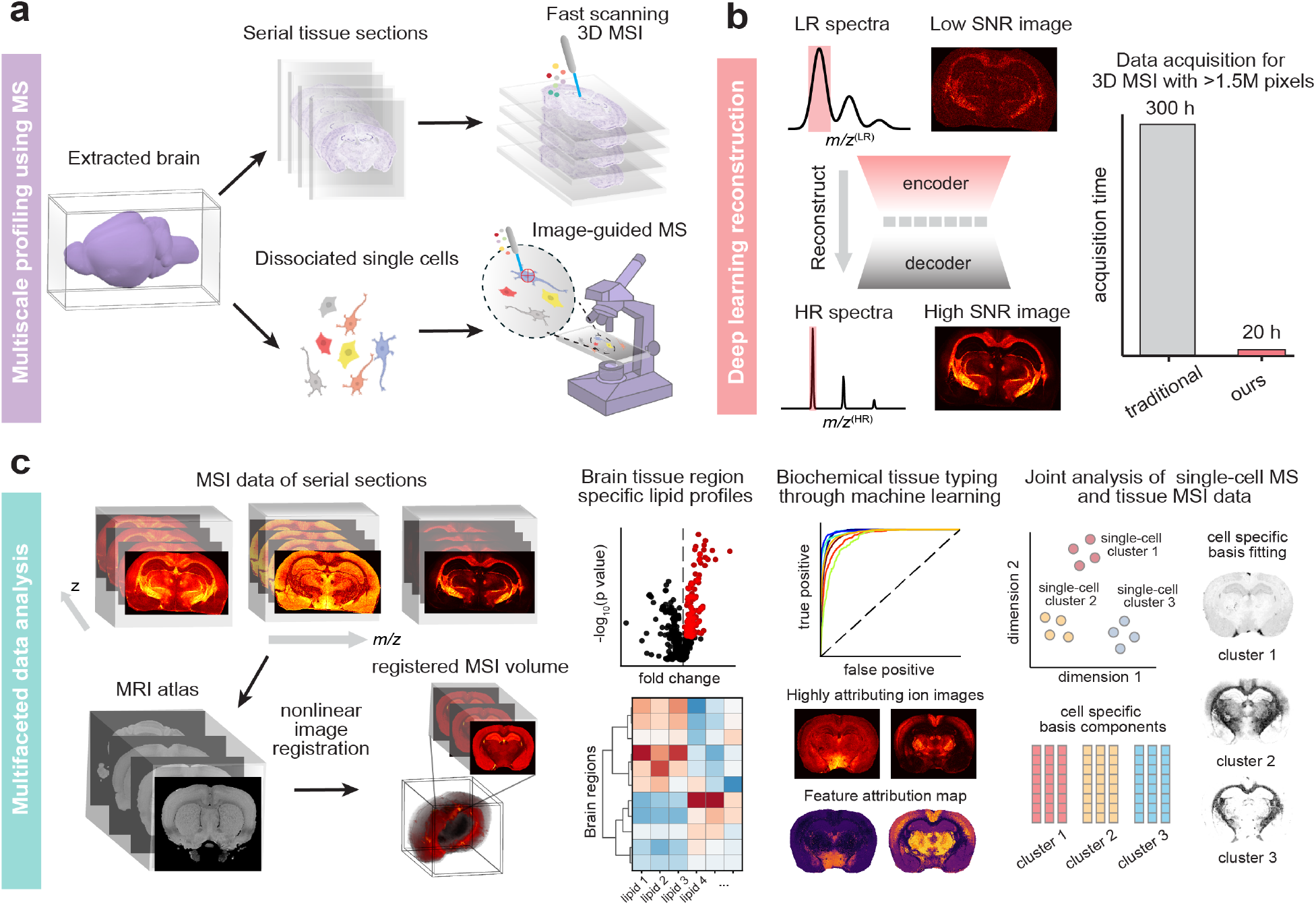
The MEISTER framework for multiscale biochemical profiling using high-mass-resolution MS enhanced by computational methods. **a**, Obtained from surgically extracted brain, serial tissue sections are imaged for 3D MSI using a fast acquisition strategy, and single cell populations prepared by tissue dissociation are probed with high-throughput image-guided MS. **b**, A deep learning model reconstructs high-mass resolving and high-SNR MS data from the low-mass resolution measurements acquired with fast acquisitions by exploiting low-dimensional manifold structure for high-dimensional MS data, producing large data sets with millions of pixels, which was previously time prohibitive with the conventional acquisition. **c**, Our multifaceted data analysis pipeline uses various data-driven methods for multimodal image registration to align MSI with 3D anatomical MRI for volumetric reconstruction, identifying differential lipid distributions, tissue typing using MS data, and integrating MSI and single-cell MS data for joint analysis and resolving cell-type-specific contributions at the tissue level across the brain.

To enable biochemical characterization of the brain and knowledge discovery from such unprecedented data, we developed and integrated several data-driven methods for analyzing the high-dimensional, multiscale 3D MSI and SCMS data (Fig. 1c). First, the MSI data were mapped to MRI and the Waxholm Space atlas through a customized nonlinear image registration procedure (Fig. 1c, left panel), which enabled a coherent volumetric MSI reconstruction from the sections imaged. Through registration, we extracted the mass spectra of 11 major brain structures across the 3D volume and identified spatially differential biochemical profiles. Next, we classified the brain structures based on their lipids for tissue typing, which identified enriched lipid species in each region/tissue type. To connect the tissue MSI and SCMS across different scales, we built cell-type-specific “chemical dictionaries” and introduced a joint union-of-subspaces (UoSS) fitting technique that resolved cell-specific contributions to the spatio-chemical contents at the brain-wide tissue level for the first time (Fig. 1c, right panel).

### Validation of the deep-learning-based MSI reconstruction

Using a carefully designed, biochemically-relevant simulated MSI dataset that contained rich chemical details and brain-mimicking spatial variations, we trained and validated the proposed deep-learning-based method for reconstructing high-mass-resolution mass spectra and ion images from noisy short transients (Extended Data Fig. 2a,b and Methods). Our method showed near ground-truth level fidelity with more than 10 dB gain in signal-to-noise ratios (SNRs) over the noisy reference using only 5% data. We compared the performance of our model for spectral and spatial feature recovery in the simulated dataset over the standard FT reconstruction and a previously described linear subspace approach^24,25^ (Extended Data Fig. 2c, d). Compared to the standard FT reconstruction from full transients, our method also yielded a higher SNR, owing to the denoising effects of the learned low-dimensional representation.

**Figure 2.**
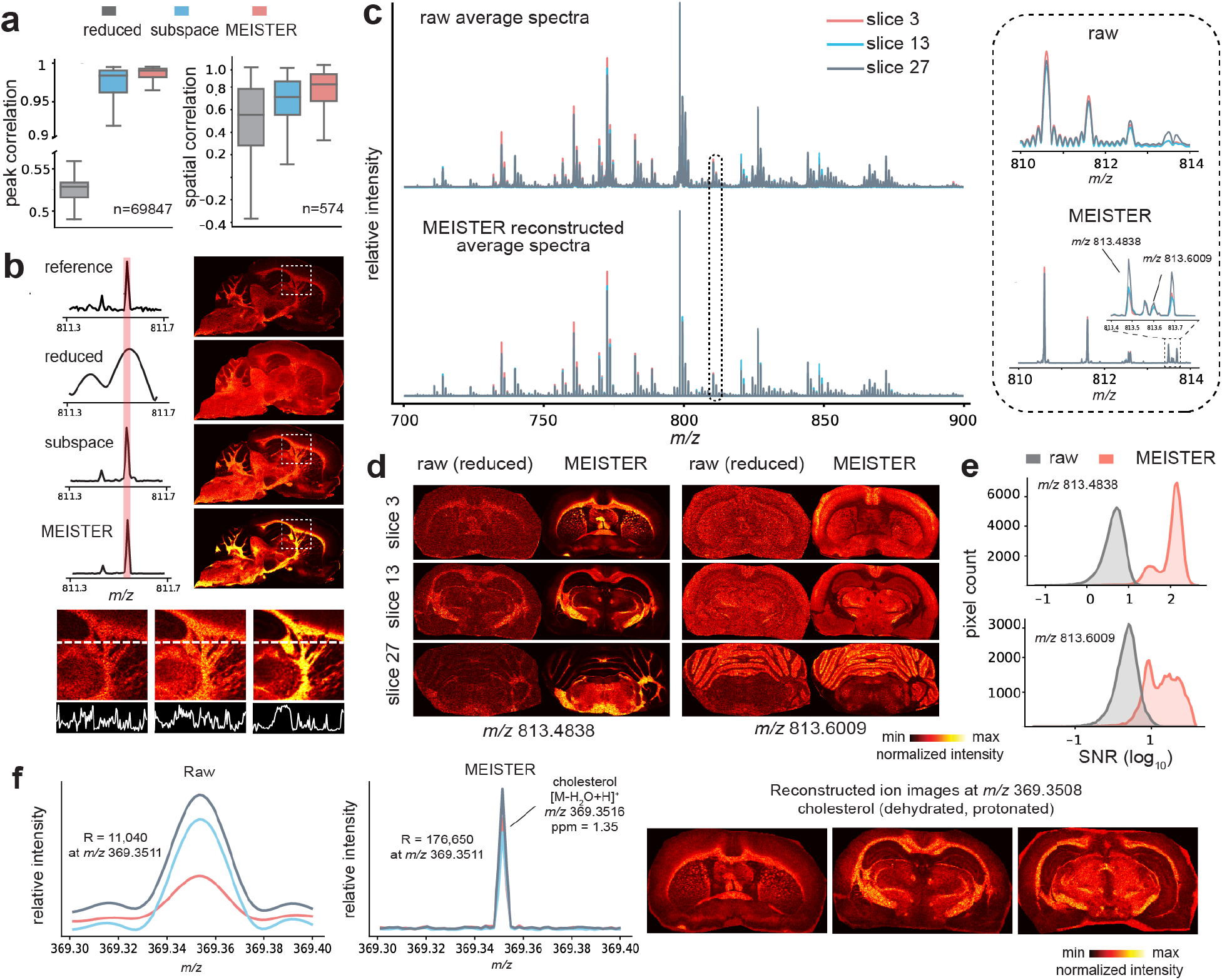
High-throughput MSI via MEISTER. **a**, Correlation measures of mass spectra produced by different reconstruction methods; the case “reduced” stands for a standard FT reconstruction from the short transients (reduced data); **b**. Comparisons of mass spectra and corresponding ion images to the reference (full transient; top row) show enhanced signal strength by MEISTER. Bottom row depicts line profiles of signal intensities in reference, subspace, and MEISTER images. **c**. Averaged mass spectra from three different tissue sections for the reduced data (top) and our reconstruction (bottom). Inlet displays a small *m/z* window of the mass spectra, demonstrating mass resolution enhancement. **d**, Ion images obtained from raw MSI data (left panel) and our reconstruction (right panel) at *m/z* 813.4838 and *m/z* 813.6009. **e**, Reconstruction improved SNRs as shown in SNR distributions for signals at *m/z* 813.4838 (top, 100-fold higher) and at *m/z* 813.6009 (bottom, 10-fold higher) than raw data. **f**. Raw and reconstructed mass spectra zoomed into the cholesterol range (left columns), and corresponding ion images showing tissue localization of cholesterol (right columns).

To evaluate our method on experimental data, we trained the model using high-resolution MSI data acquired from rat brain tissue sections using an FT-ICR mass spectrometer. We then validated the model using reference full-transient data acquired at different days from tissue sections not seen during training. For the noise contaminated reference (0.731 s transient duration, 1 million temporal points), images from peaks that were indicated in the single-pixel mass spectra showed ions with distinct spatial distributions, whereas the signals were unresolved in the reduced data with short transients (first 64,000 temporal points) due to poor mass resolution (Supplementary Fig. 1a, b). The deep-learning-based reconstruction from the reduced data successfully resolved nearby mass features, providing enhanced signal strength over the subspace-based reconstruction also from the same reduced data (Fig 2a, b, Supplementary Fig. 1b). Our method yielded quantitatively better spectral and spatial fidelity w.r.t. the reference than reduced data and subspace reconstruction. This was further supported by subsequent principal component analysis (PCA) and spatial segmentation through k-means clustering on the reconstructed spectral features (Extended Data Fig. 2e, f, g, Supplementary Fig. 2), with our method producing less noisy spatial parcellation. Our evaluation suggests that the model can learn robust nonlinear low-dimensional features from complex and noisy imaging data, while accurately predicting those features from short transients even for the highly heterogeneous brain tissue.

Furthermore, we examined the model performance for reconstructing SCMS data. Specifically, we trained a model using high-resolution SCMS data from approximately 4,000 cells, and tested on 1,000 independent cells (Methods). We found high correlation scores (Pearson r > 0.95) between full-resolution reference and reconstructed single-cell spectra. Consistent molecular profiles across individual cells (Extended Data Fig. 3a) resulted in nearly identical outcomes via uniform manifold approximation and projection (UMAP) and k-means clustering (Extended Data Fig. 3b). Even with larger chemical heterogeneity, our model was able to effectively recover the variations within and across cell populations (Extended Data Fig. 3c).

**Figure 3.**
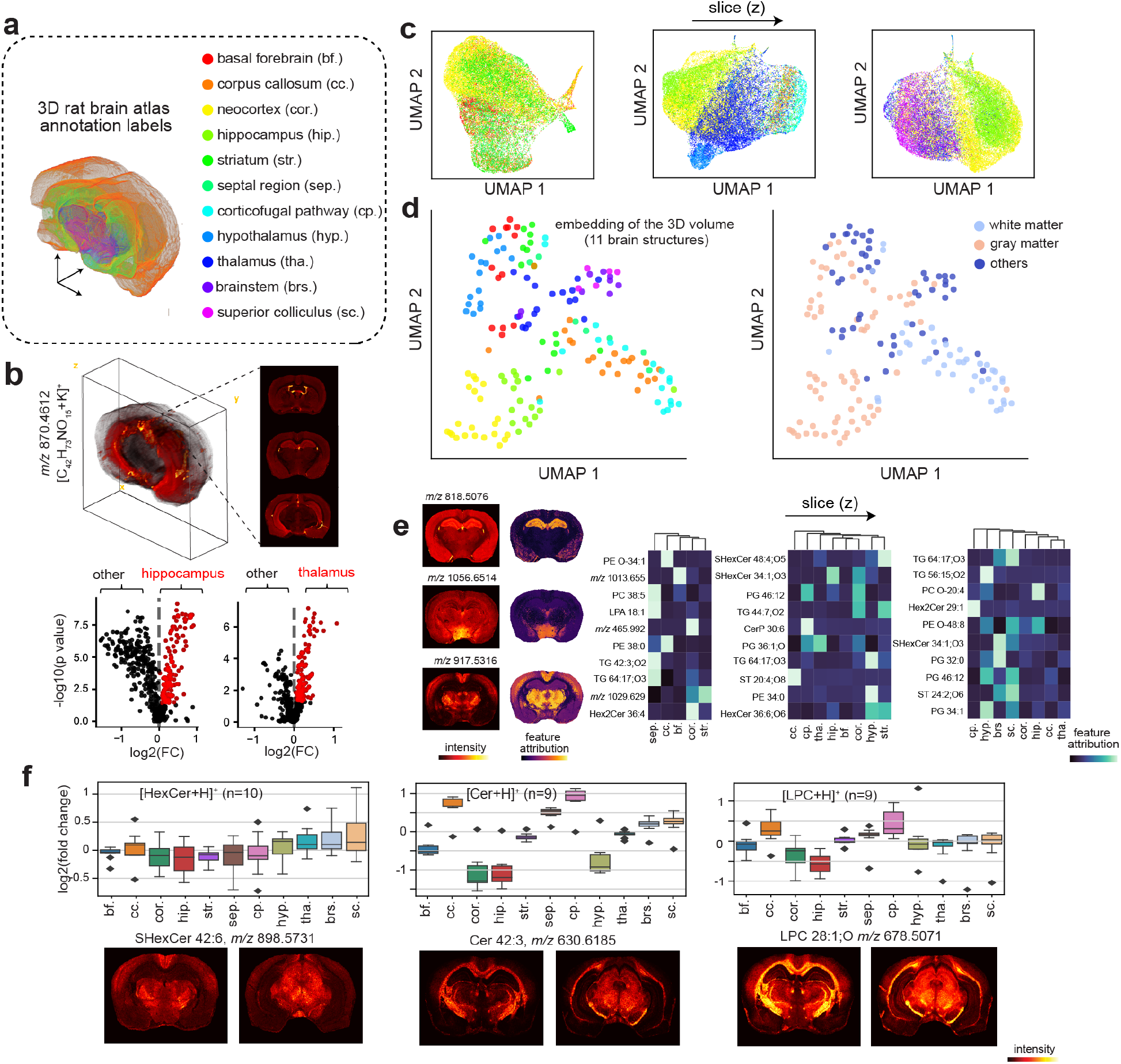
Differential lipid profiles across 11 brain structures revealed by high-resolution 3D MSI. **a**, Atlas annotations colored for 11 brain structures with abbreviations. **b**, Top: 3D volumetric reconstruction of a representative lipid ion enabled by our reconstruction and analysis methods. Bottom: volcano plots showing differential lipid distributions in hippocampus (left) and thalamus (right). Red dots indicate significantly different lipid features for the brain regions. **c**, Low-dimensional UMAP embeddings of the pixel-wise lipid profiles across different brain serial sections, revealing region-specific lipid distributions across a tissue volume. **d**, UMAP analysis of the average lipid profiles from different structures across the entire 3D volume (each dot representing a region in one tissue section). **e**, Interpreting the machine learning model trained to classify brain region using lipid profiles reveals attribution of lipids most discriminative for specific brain structures. Examples of top predictive lipid features show distinct spatial distributions and feature attribution maps across the brain (left panel). **f**, Top: regional distributions of myelination-related lipids (HexCer, Cer, and LPC) annotated protonated ion and quantified by log2 of fold change, and Bottom: representative ion images corresponding to the lipids described in the plots on top.

### High-resolution 3D MSI with large volume coverage

High-mass-resolving MSI with 3D tissue coverage has been shown^26,27^. However, the combination of mass and spatial resolution and organ coverage for volumetric imaging with FTMS has been limited, due to the inherent throughput constraint, e.g., resolution may be sacrificed (> 50 𝜇m pixel size) for imaging multiple sections for 3D imaging or the number of sections may be reduced for maintaining a small pixel size. The higher throughput afforded by MEISTER allowed us to spatially profile metabolites and lipids for many serial tissue sections that cover a large volume of the brain. To demonstrate this capability, we imaged 37 coronal and 39 sagittal sections of the rat brains, and used data with the targeted mass resolution from a few tissue sections for training (Methods). We were able to efficiently reconstruct high-mass resolution, high-SNR spectra from raw data acquired with short transient duration (i.e., <10% collection time per transient) for all remaining serial sections for approximately 2 million total pixels in each 3D data set (Extended Data Fig. 4a, Supplementary Fig. 3). Reconstructed data exhibited significantly improved quality with larger than 10-fold increment in SNRs over the raw data processed by traditional FT analysis (Fig. 2c, d, e, Extended Data Fig. 4b, c), while maintaining high mass accuracy on several expected lipid signals in rat brain and low mass errors on tentatively assigned lipids (Extended Data Fig. 4d). Our method is also applicable to different organ systems and molecules other than lipids. To demonstrate its generalizability, we imaged and trained models on rat pancreas tissue sections (Extended Data Fig 5a, b, Methods). Faithful detection and reconstruction of rat pancreatic peptides including glucagon, insulin 1 C-peptide, and insulin 2 C-peptide from reduced transient data is shown (Extended Data Fig 5c).

**Figure 4.**
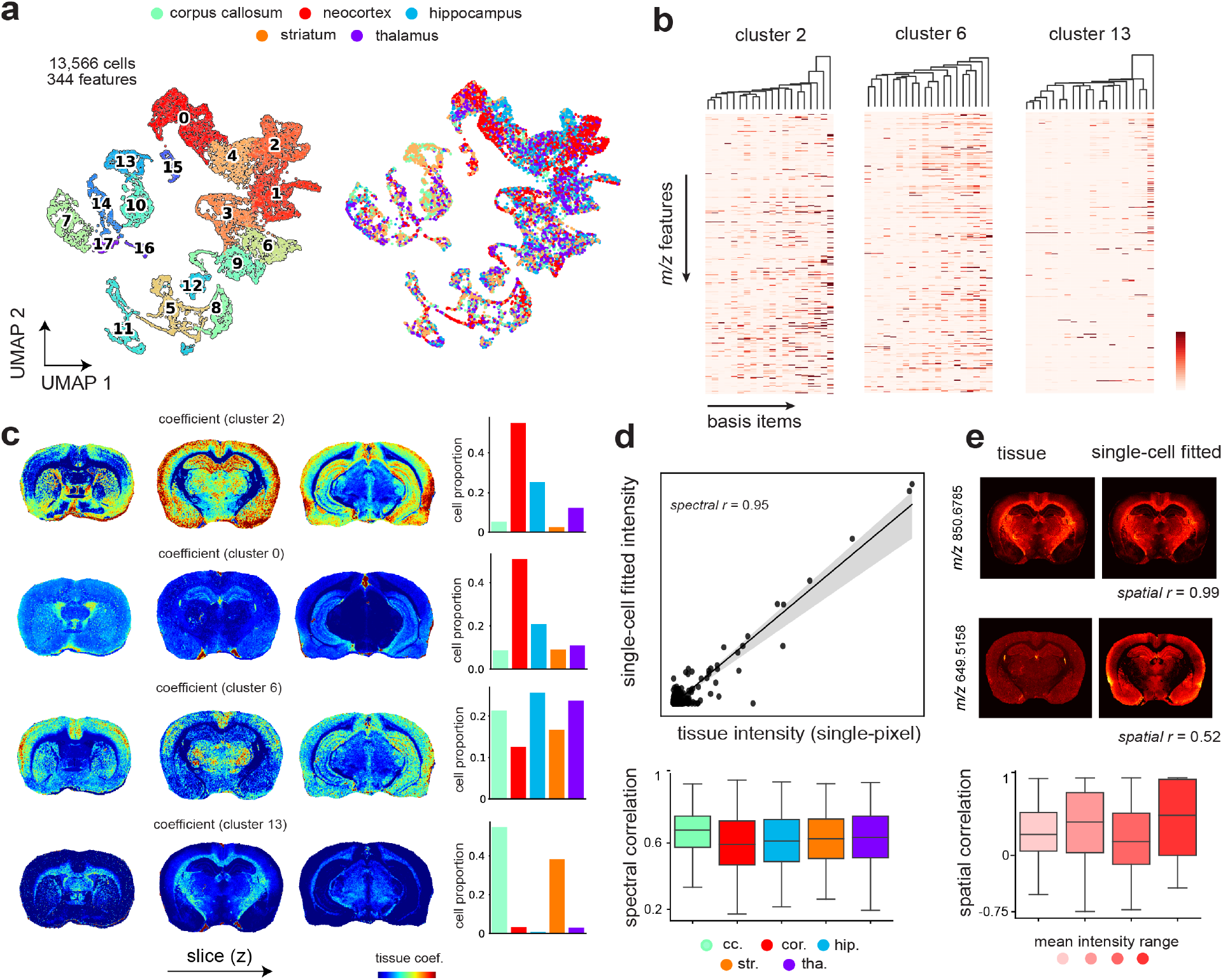
Joint visualization and analysis of tissue MSI and SCMS data. **a**, A total of 13,566 cells with 344 cross-annotated lipid features are subjected to UMAP and clustering analysis. 18 cell clusters were identified (left) and each cluster contains cells dissociated from different brain regions (right; annotated on the top). **b**, Cell-specific chemical dictionaries (lipids) can be extracted, e.g., from cluster 2, 6, and 13 shown here (20 basis elements were obtained in each dictionary). **c-e**, Results of our proposed UoSS fitting mapping cell-type-specific dictionaries (obtained from SCMS data) to the tissue MSI data. **c**, Estimated spatially dependent contributions of different cell clusters across the brain ( UoSS coefficients for the cell-type-specific dictionaries). Distinct cell compositions can be resolved for different regions. Each row shows results for mapping contributions of one cell cluster to individual pixels in different tissue sections (left three columns), and the percent compositions of regional cell populations (where they are from; right column) for each cluster. **d**. Signal intensities from tissue pixels obtained with tissue MSI are well correlated with the model fitted values (mean correlation coefficients > 0.6 for all regions, bottom box plot). **e**, Top four images illustrate lipids with excellent and moderate consistency between the actual tissue images and the single-cell-dictionary fit. The consistency is evaluated for 344 lipid features by spatial correlation across different mean signal intensity ranges. Lipids at high mean intensity have slightly better fitting results than ones at lower intensity (lower box plots). 101 (of the total 344) lipid features have negative spatial correlation, a result of less accurate fit. Brain region abbreviations: cc. corpus callosum, cor. cortex, hip. hippocampus, str. striatum, tha. thalamus.

**Figure 5.**
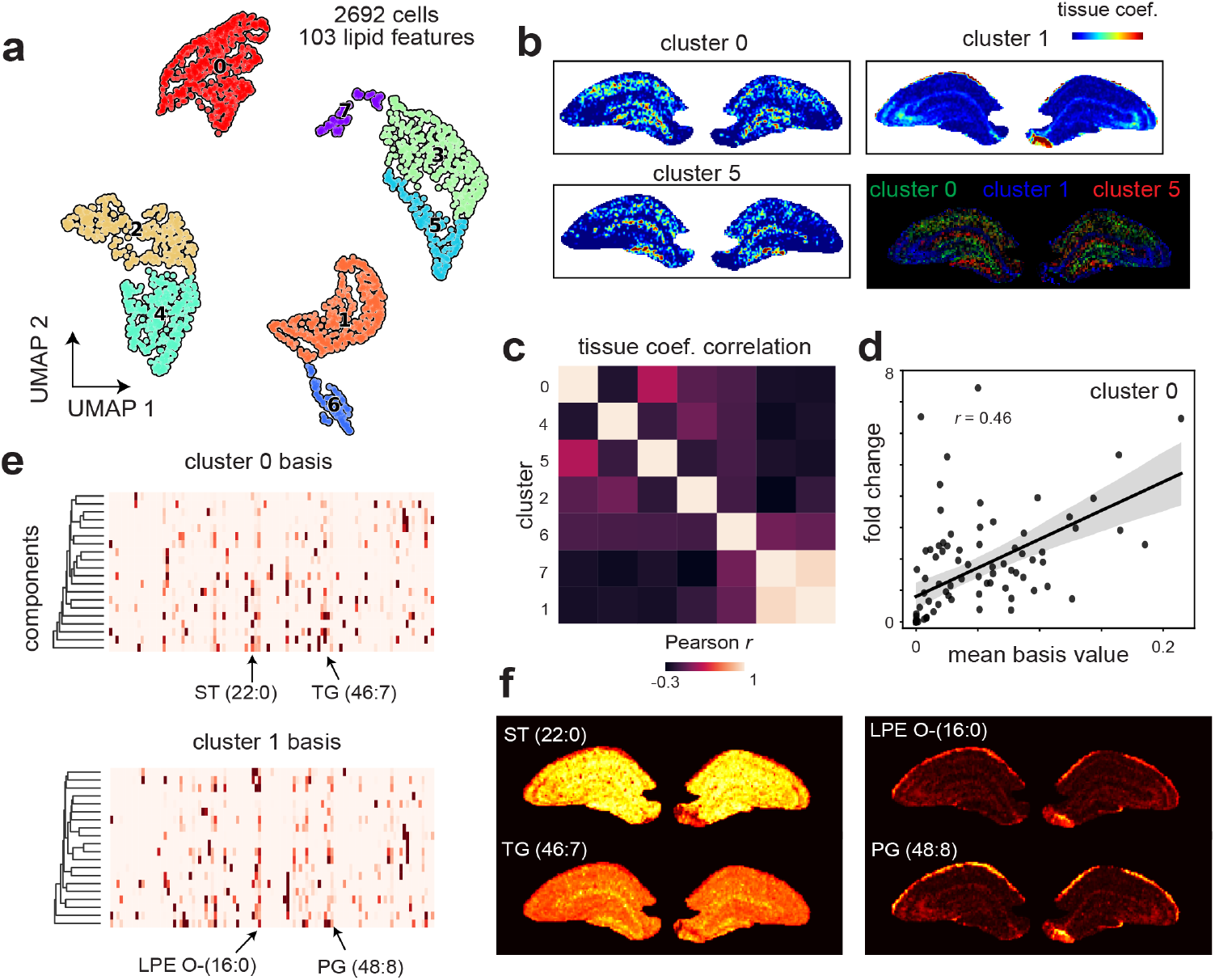
Integrative analysis of hippocampal tissue MSI and single-cell data. **a**, A total of 2692 hippocampal cells were probed and data were subjected to UMAP and clustering analysis, producing 8 chemically unique single-cell clusters in terms of lipid profiles. **b**, Maps of resolved cell-type contributions from fitting the cell-specific chemical dictionaries to tissue data. **c**, Correlation matrix for weights obtained from 8 clusters. **d**, Basis values averaged on 20 dictionary items versus fold change for lipid features in cluster 0. **e**, Chemical dictionaries extracted for cluster 0 and cluster 1, with arrows indicating dominant cluster-specific lipid features. **f**, Ion images showing the distributions of lipids that are more cluster specific within the hippocampal region. ST(22:0) and TG(46:7) were selected based on cluster 0 basis values, whereas LPE O-(16:0) and PG (48:8) were determined by cluster 1 basis.

To enable spatially resolved biochemical profiling across the brain, we designed and implemented a multimodal registration strategy to align the misaligned reconstructed MSI sections to a high-resolution rat brain MRI^28^ to form a volumetric reconstruction. Inspired by a previously proposed approach, we applied parametric UMAP^29^ to embed the MSI hyperspectral data cube into “feature” images for co-registration with MRI anatomical images (Extended Data Fig. 6a). By learning the embedding process, our method can simultaneously obtain low-dimensional representations of the entire 3D MSI data cube (Extended Data Fig. 6b, Supplementary Fig. 4 a, b). These low-dimensional features are effective in delineating tissue morphology for cross-modality image registration (Extended Data Fig. 6b, d, Supplementary Fig. 4c, d). Images of three selected UMAP dimensions of each 2D tissue section were converted to a single grayscale image to yield anatomical contrast and registered to its corresponding MRI slice via affine and B-spline registration (Extended Data Fig. 6c, Methods). The obtained transformations can then be applied to all ion images, resulting in the final high-quality volumetric reconstruction of the ion distributions (Fig. 3a, b, Extended Data Fig. 6e, f, Supplementary Fig. 5). To the best of our knowledge, this is the first time that 3D reconstruction of the biochemical distribution of the brain with the combination of coverage, spatial resolution and chemical detail (e.g., >1000 lipid features; see below) has been generated.

### 3D MSI reconstruction enabled region-specific lipid profiling across the brain

Brain lipids serve as both inter- and intracellular signaling molecules and play important functional roles in the formation of morphologically distinct membranes of the diverse neuronal cells^30,31^. Previous studies on brain lipids showed that lipid enrichments may be distinct across different brain anatomical structures^8,32,33^. With our data, a total of 1156 lipid features annotated by LIPID MAPS^34^, 728 matched with <3 ppm error were extracted from 3D MSI reconstruction for downstream analysis (Fig. 3). We performed both single-pixel analysis of individual tissue sections through UMAP visualization (Fig. 3c, Supplementary Fig. 5) and supervised classification with gradient-boosted trees. Accurate brain region classification of pixels based on lipid profiles can be achieved with an average classifier area-under-curve (AUC) of 0.96±0.02 (Supplementary Fig. 6a, Methods). We also investigated the feature attributions to interpret model decisions and selected the lipids that were contributing most to the anatomical classifications (Methods). Multivariate analysis of the 3D data was performed on the mass spectra intensity profiles of each brain region (averaged per region, per tissue section). UMAP embeddings showed a preservation of the relative spatial organizations of brain regions (Fig. 3d, see Methods for details on anatomical definitions) as well as differentiation among tissues that are gray matter dominant, white matter dominant or a mixture of both (others). Our analysis uncovered the anatomically differential lipid compositions of the brain, as shown by the ion images and feature attribution maps of the top-scoring features (Fig. 3e, Supplementary Fig. 7). The near perfect classification accuracy of anatomical structures was achieved (Supplementary Fig. 6e) using averaged region-specific lipids, indicating that MEISTER robustly uncovered anatomically specific biochemical profiles for the entire tissue volume. The mean intensities of the most discriminative lipid features from the classification model were summarized (Supplementary Fig. 6f). Among these features, we identified highly elevated region-specific lipids by comparing one structure against the rest within the tissue volume; for instance, PC O-20:4 and HexCer 40:1;O3 (PC: phosphatidylcholine, HexCer: hexosylceramide) were elevated in the hippocampal region and the corpus callosum, respectively. We further examined how sphingolipids, a lipid category critical to brain development and function, are regionally distributed (Fig. 3f). HexCer exhibits higher levels in the thalamus, brainstem, and superior colliculus, which contain large amount of nerve fiber projections responsible for sensation. Ceramide (Cer), abundant in the myelin sheath around the nerve fibers, is found elevated in subcortical white matter such as the corpus callosum and corticofugal pathway, with similar a trend observed for sphingomyelin (SM). In subcortical areas, a higher level of lysophosphatidylcholine (LPC) is also found, perhaps due to its emphasized role in myelination and neuronal membrane synthesis^35^.

### Multiscale data integration via mapping single-cell chemical profiles to tissue imaging data

Our high-throughput multiscale tissue MSI and SCMS data enabled integrative analysis to combine their powers for investigating cell-specific biochemical composition in tissues. High-throughput 3D MSI allows brain-wide biochemical characterization (e.g., lipid mapping demonstrated above), but the information at each tissue pixel contains convolved biochemical fingerprints of multiple cell types. MSI with subcellular-level resolution has been demonstrated^18,19^ but with limited tissue coverage. Furthermore, brain cells often are not organized into regular grids but interwind in complex ways. On the other hand, SCMS data acquired from individual cells dissociated from brain tissues provide cell-specific mass spectra but with limited spatial information. To integrate these two types of measurements, we reconstructed SCMS data from 13,566 cells sampled from five brain regions (same anatomical definitions as for tissue data, Methods), and annotated SCMS data using the lipid features from the tissue MSI data (considering presence of both intracellular and extracellular lipids; Methods, feature cross-annotation), resulting in 344 cross-annotated lipids in single cells. Using these lipid species, we obtained 18 single-cell clusters defined by lipid contents, with each cluster containing mixed cell populations from different anatomical regions (Fig. 4a). We characterized the distinct chemical profiles among cells across different clusters, as well as identified region-specific lipid markers, suggesting both intra- and inter-regional diversity of cellular lipids (Extended Data Fig. 7a). Single cells within the corpus callosum and the striatum contain a higher level of sphingolipids, consistent with observations from the tissue MSI data (Extended Data Fig. 7b). Differential single-cell lipid marker analysis was carried out for other brain regions, showing agreement between cellular- and tissue-level lipidomes (Extended Data Fig. 7c, d).

To integrate tissue and single-cell data and resolve cell-type-specific contributions at each image pixel, we developed a new UoSS fitting strategy exploiting cell-specific chemical dictionaries. Specifically, we performed non-negative matrix factorization (NMF) to extract sets of chemical “bases” that represent chemical variations within each cell cluster (Methods). These bases are “dictionaries” that promote sparsity and parts-based representations that delineate the biochemical components in each cell cluster^36,37^, ideal for stratifying cellular biochemical signatures. For each of the 18 clusters, we extracted 20 non-negative single-cell dictionary items (Fig. 4b, Supplementary Fig. 8b), and used a UoSS linear regression model to fit all components to tissue MSI data while forcing model weights to be non-negative (Methods). The weights can then be interpreted as the cell-type-specific contributions and yielded deconvolved cellular features at every tissue pixel (Fig. 4c). By analyzing the weights of individual cell-type clusters with respect to brain regions, we identified distinct lipid spatial organizations at the single-cell level. For example, two clusters (0 and 2) are more enriched in cortex and hippocampal regions (Fig. 4c), consistent with the observation in Fig. 4a. Although these two clusters have similar cell numbers from each region, one is more localized to the granular layer of the dentate gyrus and pyramidal layer, whereas the other is more general to Cornu Ammonis (CA) areas. Similarly, several clusters show strong spatial contributions toward the thalamus, corpus callosum and striatum (Fig. 4c). From single-cell fitting, we found a moderate to high spectral and spatial correlation of the fitted signal intensity to the original tissue signal intensity, indicating the alignment of SCMS and MSI data (Fig. 4d, e). Note that some lipid features (101 out of 344) show negative correlation between the original tissue image and the UoSS model fit using single-cell dictionaries. These might represent extracellular lipid components, false positive annotations, or modeling errors (Discussion).

To further elucidate the spatial organization of cell subpopulations within a certain anatomical region, we jointly examined hippocampal-only SCMS and tissue MSI data (Fig. 5). A total of 2,692 cells (103 annotated lipid features) were analyzed with MSI data through the joint fitting of 8 identified cell clusters (with dictionaries estimated using NMF). Single-cell lipids display heterogeneous distributions within the hippocampus, with unique lipid markers (Fig. 5a, Supplementary Fig. 9). The fitted contributions of single-cell dictionaries suggest different spatial organizations of hippocampal cell subpopulations (Fig. 5b, c). Large model weights were found in the dentate gyrus and CA3 for cluster 0, the granular layer of CA1 for cluster 1, and the molecular layer for cluster 5, respectively, approximating the morphological structure of the hippocampus (Fig. 5b). We further analyzed the extracted dictionary items, which showed strong correlation with the lipid fold change (Fig. 5d), serving as indicators of the cluster-specific lipid signatures. Features were then selected based on magnitude of the averaged basis values, which are a good proxy to lipid specificity to cell clusters. For example, consider LPE O-(16:0) and PG(48:8) (indicated in dictionary items for cluster 1, Fig. 5e). The corresponding tissue distributions for these two lipids showed agreement with the fitted model weights from cluster 1 (Fig. 5b, f).

To demonstrate the applicability of our joint analysis approach to other tissue types, SCMS data of 13,739 rat pancreatic cells consisting of islets, vasculature, and acinar cells (Methods) were acquired and reconstructed. A total 428 features were annotated using pancreas tissue MSI data. Using the cross-annotated features, we obtained 10 single-cell clusters, each containing relatively uniform cell populations (Extended Data Fig. 8a, b). Fitting the extracted single-cell dictionaries to tissue data (Extended Data Fig. 8c), we were able to map spatially dependent cell-specific contributions and resolve tissue organizations of cell populations (Extended Data Fig. 8d). For example, we observed islet populations with distinct spatial localization within the islet region (cluster 0 and 2), likely corresponding to subpopulations of islet cells.

## Discussion

We demonstrated integrative 3D tissue and single-cell biochemical mapping of the brain at a large scale using MESTER. This is enabled by a synergy of unique experimental capabilities and innovations in computational aspects including deep-learning-based reconstruction, image registration, and spatially resolved cell-specific dictionary learning and fitting. While the power of deep learning has been illustrated in various imaging modalities including MSI^38–40^, our method exploits unique signal characteristics in FTMS data and a special network design. Instead of training a deep neural network to generate high-resolution mass spectra from a low-resolution counterpart, or interpolating missing pixel values directly^41^, we jointly learn low-dimensional embeddings of the high-dimensional data and train a regression network to predict these embeddings from reduced transients, providing a strong generalization ability for both tissue and SCMS data. We validated that training can be done using different sections or animals, and works well for new datasets. Through the unique capability of MEISTER, we resolved thousands of brain lipid features over millions of pixels across a 3D volume and large cell population, while significantly reducing the data collection time. Our method should be readily adaptable to different types of molecules besides lipids and peptides (e.g., small metabolites and proteins) and other organ systems.

MEISTER relies on several alignment steps for knowledge extraction from integrating high-dimensional MSI and SCMS data. First, while alignment between MSI and other imaging modalities has been performed^42–47^, we chose to register MSI data to brain MRI which was acquired from the intact brain (without deformation) and offered a coherent 3D volumetric “atlas” for registration. We realize that the majority of prior approaches emphasized on extracting feature images from individual 2D sections, which can lead to incoherent image registration across different serial sections in large 3D MSI data sets. Our pixel-wise parametric-UMAP strategy generated feature images with a similar contrast to MRI images (Extended Data Fig. 4) and provided structurally informative features across the entire 3D data set for easier registration. The 3D registration capability can also enable richer analysis leveraging both in vivo MRI and ex vivo MSI. Second, we align SCMS and MSI data through cross-annotation to facilitate integrative analysis. Recent progress has made FDR-controlled metabolite annotation for MSI possible^48,49^. However, it is difficult to leverage such methods to annotate image-guided SCMS data due to lack of spatial information, which is a crucial statistical consideration in the annotation algorithm. Rather, we leverage the biochemical information that mutually exists in tissue and single cells, which are searched against the “tissue feature database”. This approach boosts confidence in selecting biologically viable features, as retaining mutual features can minimize the inclusion of the experiment- or sample-specific artifacts. Another advantage for cross-annotation is to consider only molecules that are present in the intracellular spaces for more accurate single-cell to tissue mapping. Meanwhile, we think it is possible to extend our approach to look at changes in extracellular spaces (e.g., in different diseases) by interrogating the fitting residual as well as lipid species filtered out via cross-annotation. For the UoSS fitting, modelling errors may occur due to falsely cross-annotated features from the tissue data caused by mass shifts and an inappropriate ppm window, or due to nonlinearities between MSI and SCMS data. These may have contributed to the poor lipid fits with negative correlation shown in Fig. 4e. A close examination of those poorly fitted lipid components may lead to important insights on potential directions for improvement in future research.

Computational methods for integrating single-cell sequencing and spatial transcriptomics (ST) data have been explored^50^, including deconvolution of cell type fractions^51–53^, joint clustering for mapping single-cell transcriptomics to ST data^54^, and estimating the number of cells per ST spot^55^. Our work is the first attempt for similar cross-scale integration for large metabolomics and lipidomics data. Our UoSS regression model is also distinct in that it does not assume only a “reference signature” but a more general mathematical representations of the ensemble chemical profiles for each cell type, capturing the intrinsic variations within each cell cluster. We used all the cross-annotated lipid species making the approach “unbiased”. In addition to identifying brain region-specific lipid variability, clustering of single cells suggests that a continuity of lipid-defined cell type diversity exists across brain regions (Fig. 4a). Similar observations have been made by transcriptomics of single cells from various brain regions, supporting that many cell types are shared between brain regions^56,57^. We have demonstrated a proof-of-concept on linking single-cell and spatial organizations of lipids, paving the way to build biochemical cartography of tissue and primary cells.

Previous studies have used liquid chromatography tandem-MS (LC-MS^2^) measurements to study differential lipid contents within different brain regions and cell types^8,32^. Comparing the LC-MS^2^-based shotgun lipidomics and our MS imaging-based lipid profiling, we have found agreements in region-specific distributions of major lipid classes, including HexCer, Cer, and SM that are more enriched in regions such as the corpus callosum with higher myelin content^32^ (Fig. 3d, e). Our multiscale imaging-based approach offers not only the capability of resolving hundreds of lipid molecules, but also a new tool for understanding spatial-biochemical tissue architecture with cellular specificity, transforming how we study brain chemistry just as how spatial transcriptomics transforms determination of gene expression. We envision future endeavors on creating multiscale biochemical atlases, with increasingly powerful profiling technology for metabolites, lipids, peptides and proteins, as well as integrative analysis with other omics data.

## Online Methods

### Experimental Details

#### Animals

The male Sprague-Dawley® outbred rats (*Rattus norvegicus*) used in this study were sourced from Inotivco (www.inotivco.com). These rats were provided with ad libitum access to food and housed on a 12-h light cycle. All animal euthanasia procedures were carried out in strict accordance with the guidelines set forth by the Illinois Institutional Animal Care and Use Committee as well as the federal and ARRIVE guidelines to ensure the humane care and treatment of animals.

#### Tissue Dissociation and Preparation of Single Cells

A total of 3, 2.0–2.5-month-old male rats were used for brain tissue isolation. Each isolated tissue region was individually treated with a papain dissociation system (Worthington Biochemical, Lakewood, NJ) and incubated for 120 min at 34 °C with oxygenation. The treated tissue regions were then mechanically dissociated in ice cold modified Gey’s balanced salt solution (mGBSS) containing (in mM): 1.5 CaCl_2_, 5 KCl, 0.2 KH_2_PO_4_, 11 MgCl_2_, 0.3 MgSO_4_, 138 NaCl, 28 NaHCO_3_, 0.8 Na_2_HPO4, and 25 HEPES, pH 7.2, and supplemented with 0.08% paraformaldehyde to stabilize cells against damage during dissociation and other methodological steps. A solution of 80% glycerol and 20% mGBSS was added to a final glycerol concentration of 40% (v/v). The cells were stained with Hoechst 33342 (0.1 μg/mL in mGBSS) and a 30 μL aliquot of cell suspension was plated onto an indium tin oxide (ITO)-unpolished float glass slide, Rs = 70-100 Ω (Delta Technologies, Loveland, CO). After ∼16 h, glycerol was aspirated off the dissociated cells before rinsing with 150 mM ammonium acetate. Each slide held 3 biological replicates of each brain region and were placed in discrete but random locations on the ITO-glass slide to mitigate batch and spatial-dependent artifacts.

Islet isolation was performed as previously described^58,59^ with some modifications. Briefly, the pancreas was surgically removed and treated with lyberase. Islets were manually collected by mechanical dissociation of tissue using a micropipette under visual control using an inverted microscope. Islets, acinar tissue and vasculature regions were incubated for 20 minutes at 37 °C in Trypsin LE solution before mechanical dissociation into single cells and deposition onto ITO-coated glass slides.

#### Tissue Sectioning

Coronal and sagittal brain slices were obtained from the rats in this study. The entire rat brain was quickly removed and flash-frozen after decapitation, before being sectioned. Sagittal brain slices were prepared at a temperature of –20 °C and sliced into 16-μm-thick tissue sections using a cryostat-microtome (3050S, Leica Biosystems Inc., Buffalo Grove, IL). The tissue slices were then thaw-mounted onto ITO-coated glass slides for MALDI matrix application. The pancreas from 3 male rats were surgically removed and immediately frozen. Six 16 µm thick adjacent sections were cut from the frozen pancreas using a cryostat-microtome and similarly thaw-mounted onto ITO-coated glass slides for matrix application.

#### Matrix Application

The MALDI matrix 2,5-dihydroxybenzoic acid (DHB) was prepared to a concentration of 30 mg/mL in 70% methanol for brain samples. For pancreatic samples, DHB was prepared at a concentration of 10 mg/mL in 50% ethanol. Matrix was applied using an HTX-M5 Sprayer (HTX Technologies, Chapel Hill, NC), with a spray spacing of 2.5 mm at a temperature of 75 °C using a flow rate of 100 μL/min. The distance of the sprayer nozzle was 50 mm from the sample and a spray pressure of 10 psi with a spray nozzle motion velocity of 1200 mm/min was used. Four passes were used to apply MALDI matrix.

#### Image Guided SCMS Analysis

The Brightfield and fluorescent microscopy images were obtained using an Axio Imager M2 (Zeiss, Jena, Germany) equipped with an AxioCam ICc 5 camera and a .63× camera adaptor. For transmitted light, a VIS-LED lamp was used, while for fluorescence, an X-cite Series 120 Q mercury lamp (Lumen Dynamics, Mississauga, Canada) was employed. The imaging was done using DAPI (ex. 335–383 nm; em. 420–470 nm) dichroic filter cubes. The images were acquired in mosaic mode with a 10× objective and 10% tile overlap. The resulting tiles were stitched together before being exported in TIFF-file format using ZEN 2.0 Pro edition (Zeiss, Jena, Germany) software. The single-cell coordinates, geometry files, and an Excel file required for the target automation function on ftmsControl (v. 2.1.0, Bruker Corp., Billerica, MA) were obtained using microMS, as described previously^23^. To ensure data quality, cells were filtered from lists of analyzed structures based on their distance from each other, with cells closer than 200 μm being removed, and based on their size, with any free nuclei resulting from cell lysis being removed. High-throughput single-cell analysis was performed using a SolariX 7T FT-ICR mass spectrometer (Bruker Corp.), with a mass window of 50–1000 *m/z* (rat brain) or 150-6000 *m/z* (rat pancreas). MALDI mass spectra were acquired in positive mode using a Smartbeam-II UV laser in "ultra" mode, which produces a 100 μm diameter laser-spot size. Each MALDI acquisition was comprised of one ICR accumulation, consisting of 150 or 500 laser shots, for brain and pancreatic samples respectively, at a frequency of 1000 Hz.

#### Signal modeling

A transient can be modeled as a temporal signal that contains many frequencies corresponding to different ions, following the generic signal model proposed by Marshall^60,61^:

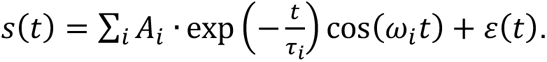

For ion 𝑖, constant 𝐴_i_, represents the initial signal amplitude, 𝜏_i_, is the decay rate of the excited ICR signal due to ion collisions, 𝜔_i_, is the ion cyclotron frequency, and 𝜀 is the independent noise. The theoretical mass resolution is calculated as:

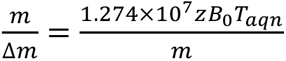

where 𝐵_0_ is the magnetic field strength, and 𝑇_aqn_ is the transient acquisition time. Given a fixed 𝐵_0_, the theoretical mass resolution is thus proportional to 𝑇_aqn_: a certain 𝑇_aqn_ is required for a target high mass resolution. We used the described signal model for the MSI data simulation.

#### Simulation of MSI data

We followed a previously described procedure to simulate the MSI data^25^. Briefly, transients were generated through the generic signal model discussed above with a list of 30 chemical formulas. Frequencies were reverse calculated for all possible ions including H+, Na+, and K+ adducts were assigned to each formula, and theoretical isotopic distributions were calculated using the Python version of the BRAIN algorithm^62^. Allen Brain Atlas (ABA) mouse brain annotation was used as the spatial reference^63^ to generate 8 pseudo-tissue regions with different combinations of chemical formulas. All transients were simulated for 262,144 temporal data points in total of 26,497 pixels. Independent Gaussian noise was added to each simulated transient.

### Model design for MEISTER

#### Reconstruction model

The signal reconstruction model consists of three parts: 1) an encoder network encoding input high-resolution transient signals into lower-dimensional latent features, 2) a regression network transforming corresponding low-resolution signals to their latent features, and 3) a decoder network decoding the estimated low-dimensional latents back to high-resolution signals. We use a deep autoencoder architecture to learn low-dimensional features directly from raw high-resolution transient signals for both tissue and single-cell measurements. Each transient signal 𝑠(𝑟*_n_*, 𝑡) is sampled with a specific temporal sampling rate, with 𝑡 = {𝑡_1_, 𝑡_2_, … 𝑡*_NT_* } where 𝑁*_T_* is the number of discrete time points and the duration 𝑇 for a defined mass resolution, and 𝑛 = 1,2, ⋯ , 𝑁_r_, where 𝑁_r_ corresponds to number of pixels in MSI or number of cells in SCMS data. Denoting 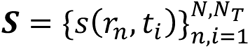 as the ensemble of training data, our objective is to train the network to encode 𝑺 into a set of low-dimensional features and produce reconstruction 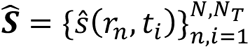 that are as close to 𝑺 as possible. Specifically, our network can be described mathematically as:

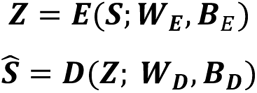

where 𝒁 represents a 32-dimensional latent vector encoding 𝑺, 𝑬(⋅) **and** 𝑫(⋅) denote the encoder and decoder functions, respectively, each containing three fully-connected layers with 512, 256, and 64 neurons respectively (symmetric design). Denoting the whole network as 𝜙(⋅; 𝚯) (combining encoder and decoder) with 𝚯 = [𝑾_𝑬_, 𝑩_𝑬_, 𝑾_𝑫_, 𝑩_𝑫_] containing all the network parameters, the MSE loss was minimized during training:

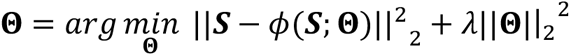

We then trained a regressor network 𝑹(⋅) to map the low-resolution measurements 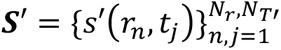, with 𝑁_*r*_ transients (𝑁_*r*_ ≫ 𝑁) and first 𝑁*_T_*, temporal points corresponding to a shorter acquisition duration 𝑇′ (𝑇′ ≪ 𝑇), to the latent features 𝒁′:

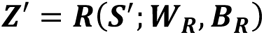

which can be decoded into full-resolution transients. Denoting all the regressor network parameters as 𝚯_𝐑_ = [𝑾_𝑹_, 𝑩_𝑹_], the model was trained by minimizing the mean-squared-error between 𝒁 (encoded from 𝑺) and the regressor output 𝒁′:

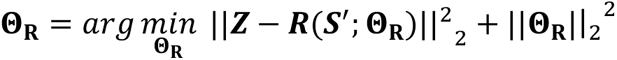

Low-resolution measurements 𝑺′ can then be transformed into high-resolution data by:

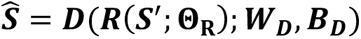

#### Evaluation on simulated and experimental MSI data

First, we validated MEISTER’s signal reconstruction performance on simulated MSI data. We trained MEISTER on 2,000 randomly sampled pixels containing noisy transients (simulation data described above), and performed reconstruction on all pixels containing reduced noisy transients (taking the first 10,000 temporal points). We compared the reconstructed data against the ground truth high-resolution data without Gaussian noise. All sets of transients (clean, noisy, noisy reduced, and reconstructed) were transformed into mass spectra and converted to peak intensity lists, which consist of peak centroids identified from the ground truth spectra. Peak and spatial correlation scores against the ground truth were computed as the Pearson correlation coefficients between each peak list and ion image pairs. Encoded features of the latent space were extracted from the bottleneck layer and subjected to UMAP for visualization.

For the experimental data, we trained the networks using a set of high-resolution data (1M temporal points) acquired on rat sagittal and coronal brain sections and the corresponding low-resolution data (64K temporal points). Reconstruction was performed on the 160 µm, 1 mm and 2 mm sections away from the training sections to validate the model’s generalizability across tissue volume. High-resolution data were acquired from these tissue sections (serving as the reference) and reduced to 64K temporal points as the input to MEISTER reconstruction model. Reduced (zero-padded), reconstructed and reference transients were then transformed into mass spectra and converted to peak intensity lists. For each tissue section, the peak centroids were determined on the average mass spectra obtained from the high-resolution reference data. Peak and spatial correlation scores were calculated the same way as for the simulation, but against the high-resolution reference. SNRs were defined as the ratios between the signal intensity and the standard deviation of the noise, which was obtained over a spectral region without apparent signals. K-means clustering was performed for different reconstructions with k=6. The components and scores for first 5 PCs were compared between reconstructed and reference data.

#### MEISTER for 3D MSI

For 3D MSI of rat coronal sections, training data (transients collected for 1M temporal points) were collected on 3 tissue sections with a total number of 124,370 pixels. For rat sagittal sections, data from 2 tissue sections in total of 105,954 pixels were used for the model training. Training sections were roughly 2 mm apart to ensure the coverage of diverse tissue types. The autoencoders were trained for 20 epochs and the regressors were trained for 50 epochs. A batch size of 128 and Adam optimizer were applied to train both networks. We then acquired low-resolution data with 64K temporal points (mass resolution 10,000 at *m/z* 400) for all remaining tissue sections (37 coronal and 39 sagittal). During reconstruction, these low-resolution signals were served as input for the regressor network, which predicted 32-dimensional latent vectors for each signal. The predicted latent vectors were decoded to transient signals with 1M temporal points (mass resolution 160,000 at *m/z* 400) by the previously trained decoder. Finally, high-resolution mass spectra were generated from the decoded transients.

#### MEISTER for image-guided SCMS

We trained MEISTER using 3,840 random cells (data with 1M temporal points) from five brain regions (neocortex, hippocampus, thalamus, striatum, corpus callosum) using microMS. The autoencoder and regressor were trained for 20 and 50 epochs respectively, with a 64 batch size. Model was validated on a validation set containing 1,000 cells. The spectral correlation scores were calculated between the reference peak intensity and the reconstructed peak intensity. To show MEISTER reconstruction provides consistent downstream analysis, we compared UMAP and Leiden clustering results between the reference (validation set) and reconstructed single-cell data, and visualized the single-cell distributions of *m/z* features over UMAP (Extended Data Fig. 3). A total number of 13,566 cells (64K temporal points) was reconstructed (1M temporal points) to obtain high-resolution single-cell mass spectra.

### Analysis of 3D MSI data

#### Data preparation

To prepare for data analysis, we first determined the peak centroids on the average mass spectra for each tissue section, and extracted the intensities of the peak centroids for all pixels. The peak lists were then processed by *m/z* binning in 3 ppm increments to align peaks affected by potential mass shift. After extracting peak lists from a 3D data set, we retained *m/z* bins common across all tissue sections for further analysis. Each pixel was normalized by total ion count (TIC). The processed data were finally converted into imzML file format.

#### Data-driven image registration

To enable 3D reconstruction and analysis of MSI data with respect to brain anatomy, we registered MSI serial sections to T_2_*-weighted anatomical MRI from Waxholm Space atlas of the Sprague Dawley rat brain. To ensure precise and accurate registration across serial sections, we adapted parametric-UMAP to extract both structurally informative and consistent feature images from high-dimensional MSI data. Previous work has demonstrated using low-dimensional feature images (embeddings) obtained via nonparametric dimensionality reduction methods (both t-SNE and UMAP) for image registration tasks. However, feature images from different tissue sections can provide disparate morphological contrasts, since data are essentially embedded into different embedding space per tissue section. Embedding the entire 3D MSI data set can overcome such issue, but it is computationally intractable for t-SNE and UMAP optimization over millions of input pixels with thousands of dimensions. The major advantage of parametric version of UMAP is to use a neural network to learn a relationship between data and embedding. Thus, a small subset of pixels can be sampled from 3D MSI data for training the network, which can rapidly embed a large number of pixels into a single embedding space. We used an autoencoder in conjunction with UMAP, of which the encoder is trained to minimize UMAP loss and the decoder is trained to minimize reconstruction loss. The autoencoder input and output size is set to be the number of *m/z* features, followed by 256, 128, 64 size of fully connected layers. The UMAP loss function between two data points 𝑖 and 𝑗 is the cross-entropy defined as:

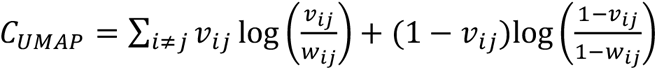

Ten percent of total pixels were randomly sampled as training data, and the network was trained for 20 epochs. For each tissue section, embedding vectors from three UMAP dimensions were encoded into RGB channels to form feature images, which were then converted to grayscale ready for registration.

The anatomical images from MRI atlas were selected based on the tissue sectioning distance with manual inspection. We applied a two-step multimodal image registration to align grayscale MSI feature images as the moving images with the reference anatomical images. First, rigid affine registration was performed to roughly align the two with 9 hand-selected initial transformation points. After rough alignment, a nonrigid cubic B-spline registration was performed with Mutual Information as the similarity measure with 200 maximum optimization steps. The registration quality was evaluated via Dice Index (DI), which assesses the image mask overlap between the 𝑖-th brain region labels from the atlas and the human-annotated masks from the 𝑗-th registered tissue section (Supplementary Fig. 10):

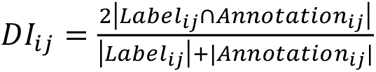

The transformation matrices were applied to each MSI section to visualize registered ion images. The regional-specific mass spectral profiles were extracted from pixels based on the atlas brain region labels.

#### Brain regional-specific analysis

To analyze regional lipid distributions at the single-pixel level, we extracted all MSI pixels belonging to 11 brain structures based on an MRI atlas labels, including neocortex, hippocampus, thalamus, hypothalamus as gray matter dominant regions corpus callosum, corticofugal pathway as white matter dominant regions, and superior colliculus, basal forebrain, brainstem, striatum, and septal regions as regions (others) that contain collections of clusters of cell bodies as well as processes for signal communication. For individual tissue section, the extracted pixels were first subjected to UMAP for visualization and used to train Gradient boosting tree (GBT) models to predict the brain regions, a multiclass classification task. Training and testing set sizes were set to 0.8 and 0.2. GBT models were further interpreted through SHAP (SHapley Additive exPlanations) values. In SHAP, each pixel provides the lipid feature attributions toward predicting certain brain regions, which can be used to generate feature attribution maps for ion images. The most contributing lipid features were selected by ranking mean absolute SHAP values. Regional average lipid profiles were obtained from every tissue section, which were repeated for the aforementioned analysis. Differential analysis of lipid features was performed for each brain region to obtain the log2 fold-change and p-values tested by Wilcoxon rank-sum and adjusted by Benjamini-Hochberg. For putative lipid annotation, we searched the *m/z* values against LIPID MAPS^34^ experimental and virtual databases with a +-0.005 *m/z* threshold for chemical formula and lipid species assignments. From the combined list, the matches were sorted according to their ppm errors from the accurate masses. In cases when experimentally or structurally validated lipids (biologically relevant lipids present in LMSD) are matched, they are given priority for assignment.

### Joint analysis of MSI and SCMS data

#### Cross annotation strategy

We applied a straightforward strategy to annotate lipids in SCMS data using features observed in MSI data. Similar to putative annotation, features in MSI were served as a database to search the SCMS peak lists for matching lipids within a 3 ppm *m/z* window. Features present in less than 5% of cells were discarded, and cells with less than 5% total number of features were filtered out. Using an alternative method, we first annotated the tissue MSI data using METASPACE with CoreMetabolome database and obtained the monoisotopic *m/z* features from the annotation results with a 50% FDR. These monoisotopic ions were then used to search the SCMS data.

#### Integrative analysis using union-of-subspace fitting

The 𝑚x𝑛 single-cell lipid feature matrix 𝑿 were first processed by TIC normalization, with 𝑚 being the number of cells in each identified cluster and 𝑛 being the number lipid features. Leiden clustering was performed with on the first 40 PCs with the parameters n_neighbors=30, min_dist=0.5, and resolution=0.25, and cosine as the distance metric. The single-cell matrix 𝑿^(𝒍)^ of cell cluster 𝑙 can be decomposed into:

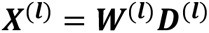

where 𝑾^(𝒍)^ represents the 𝑚x𝑘 weight matrix (𝑘 being the number of dictionary item capturing the chemical variations within each cluster), and 𝑫^(𝒍)^ is the 𝑘x𝑛 nonnegative dictionary matrix that contains sparse representations of lipid signatures in 𝑙-th single-cell cluster. We choose 𝑘=20 for the NMF algorithm. The union of dictionary items concatenated across 𝐿 number of clusters 𝑼 = [𝑫^(𝟎)^; 𝑫^(𝟏)^; … 𝑫^(𝑳)^] is fitted to the 𝑝x𝑛 tissue imaging data matrix 𝒀, where p being the number of tissue pixels, by a constrained linear least-squares fitting, with model weights constrained to be nonnegative:

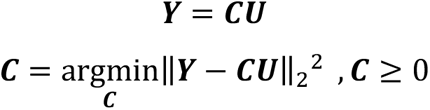

The 𝑝x𝑘*𝐿 tissue weight matrix 𝑪 has the row vectors 𝑪*_pi_*, = [𝑐*_pi_*_,0_, 𝑐*_pi_*_,1_ … 𝑐*_pi,L_*] and 𝑐*_pi,l_* contains the weights for pixel 𝑝*_i_*,and 𝑘 dictionary items from the 𝑙-th single-cell cluster. The vector norm |𝑐*_pi,l_*|_2_ was taken corresponding to the summarized contributions from all dictionary items of the 𝑙-th cluster. The vector norms were mapped back to original pixel locations to visualize the spatial contributions of single-cell lipid signatures at the tissue level.

## Data availability

The processed 3D MSI, single-cell MS and other relevant imaging data that support the findings of this study are publicly available and free to download from Figshare upon publication of the work. Due to large file sizes, raw data including simulated and experimental transients can be available upon reasonable request to the corresponding authors to arrange data sharing.

## Code availability

The code used in this study are free for noncommercial use and available on GitHub (https://github.com/richardxie1119/MEISTER).

## Acknowledgments

This project was supported by the National Institute on Drug Abuse under Award No. P30 DA018310, the National Institute on Aging under Award No. 1R01AG078797, the National Human Genome Research Institute under award No. RM1HG010023, and the National Institute of General Medical Sciences under award No. 1R35GM142969. The content is solely the responsibility of the authors and does not necessarily represent the official views of the awarding agencies.

## Author contributions

Y.R.X, F.L. and J.V.S. conceived the project idea. Y.R.X. and F.L. developed the deep learning and computational methods. S.S.R. and D.C.C. designed and performed the sample preparation for tissues and single cells. Y.R.X., T.J.T., and D.C.C. acquired mass spectrometry data. Y.R.X. implemented and evaluated the computational algorithms. Y.R.X, T.J.T., F.L. and J.V.S performed data analysis. Y.R.X. created the figures and all authors wrote the manuscript. F.L. and J.V.S. acquired funding and provided direction throughout the project.

## Additional information

Supplementary information is available online.

## Extended Data Figures

**Extended Data Fig. 1.**
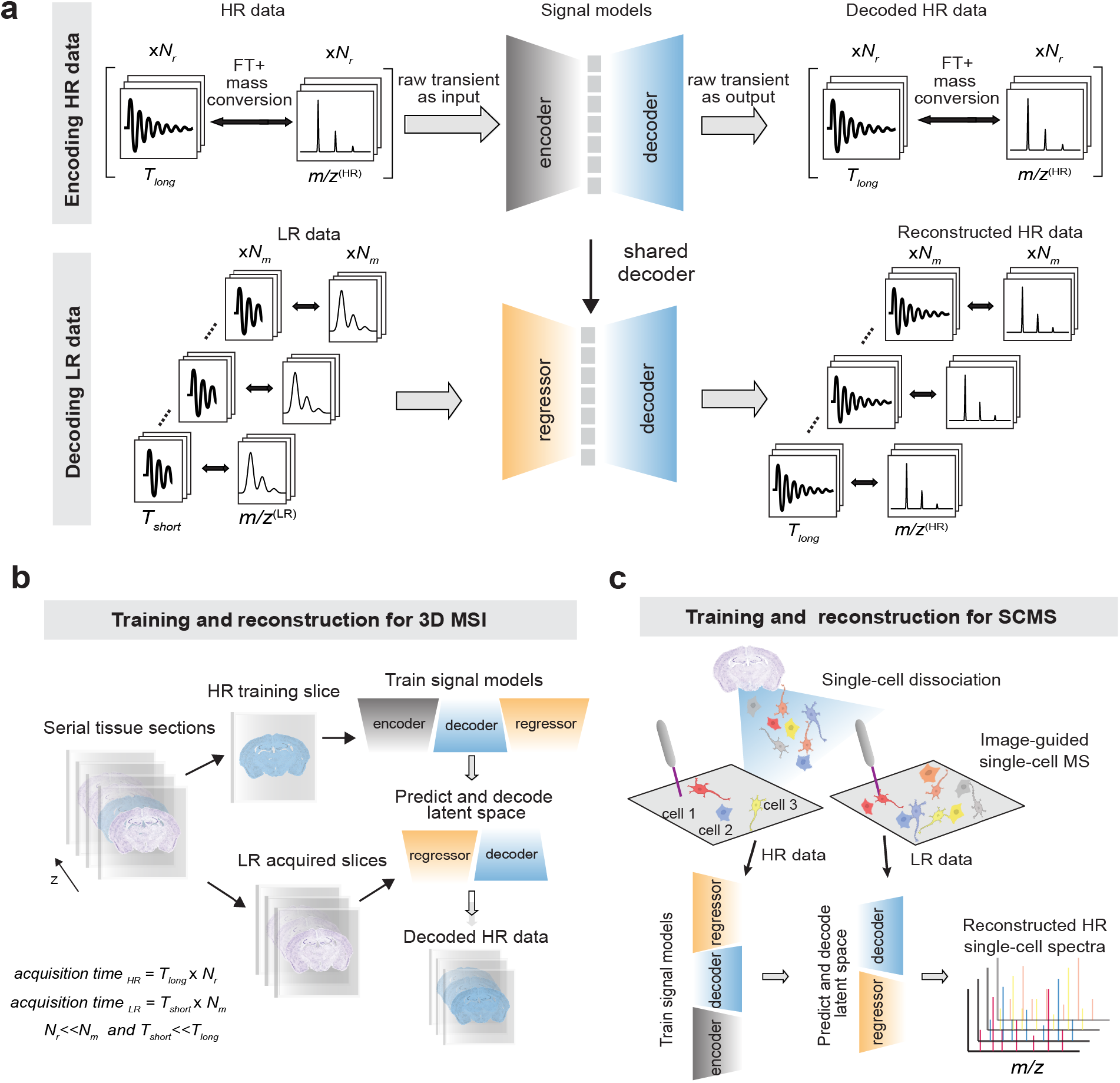
**a**, MEISTER reconstruction model that contains an autoencoder to learn latent features from high-resolution signals, and a regressor network that maps low-resolution signals to encoded latent features. MEISTER training workflow for **b,** 3D MSI using high-mass resolution data acquired on a small number of tissue sections, and **c,** SCMS using high-mass resolution data acquired on a subset of individual cells.

**Extended Data Fig. 2.**
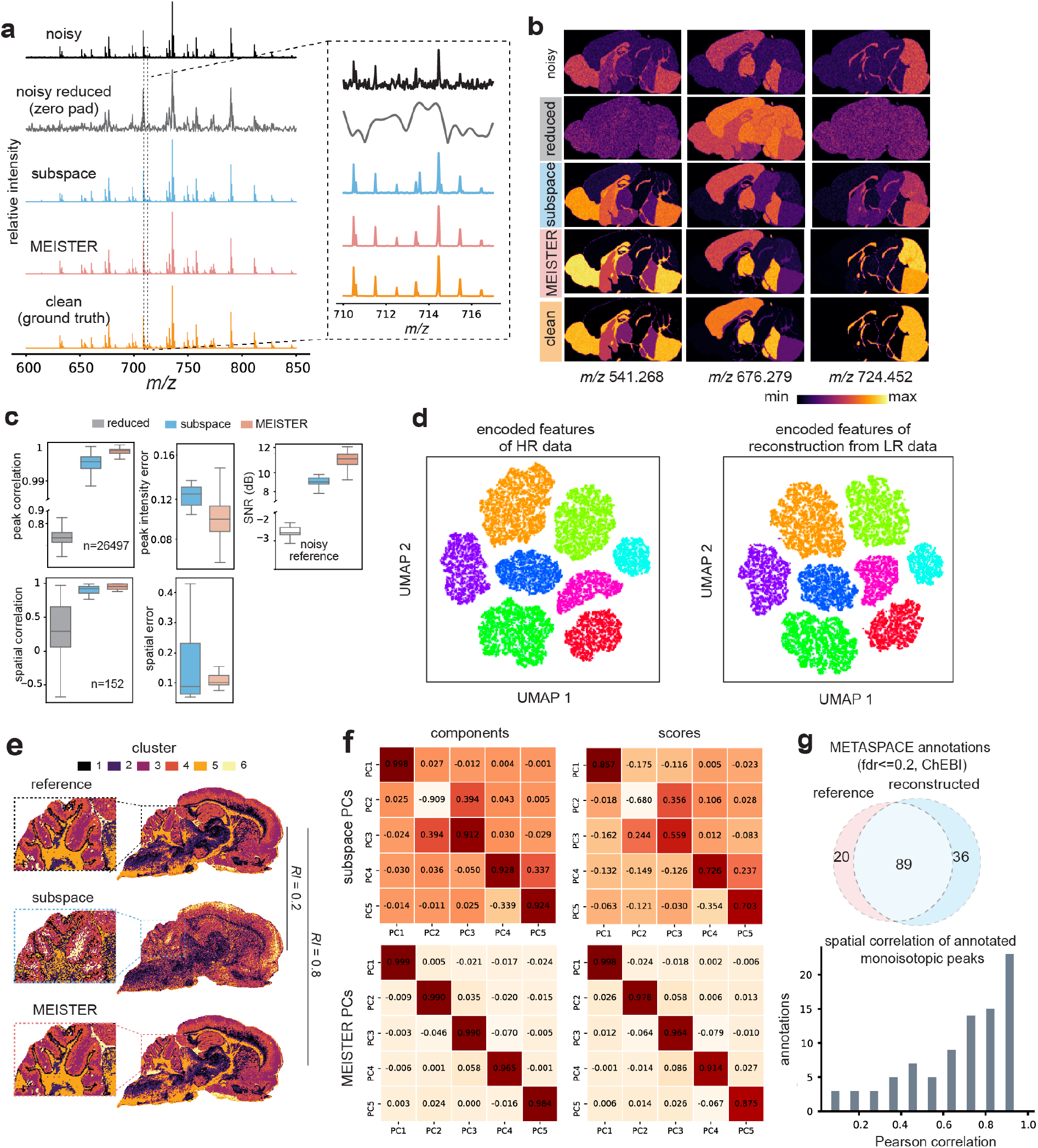
Comparisons of **a,** mass spectra from the simulated MSI data. **b,** ion images extracted from several *m/z* features, showing enhanced spectral and image quality enabled by MEISTER reconstruction. **c,** Correlation coefficient and error distributions by evaluating mass spectra and ion images against the ground truth. **d,** UMAP embeddings of encoded features of the simulated high-resolution data and the features of reconstruction from low-resolution data. Colors indicate different pseudo-tissue regions. **e,** K-means clustering for the experimental reference, subspace reconstruction, and MEISTER reconstruction. **f,** Pearson correlation coefficients between top-5 PCs extracted from the experimental reference versus from the data reconstructed by subspace (top) and MEISTER (bottom). **g**. Comparison of number of annotated lipids (top) and correlation of ion images (bottom) using METASPACE with FDR set to 20%.

**Extended Data Fig. 3.**
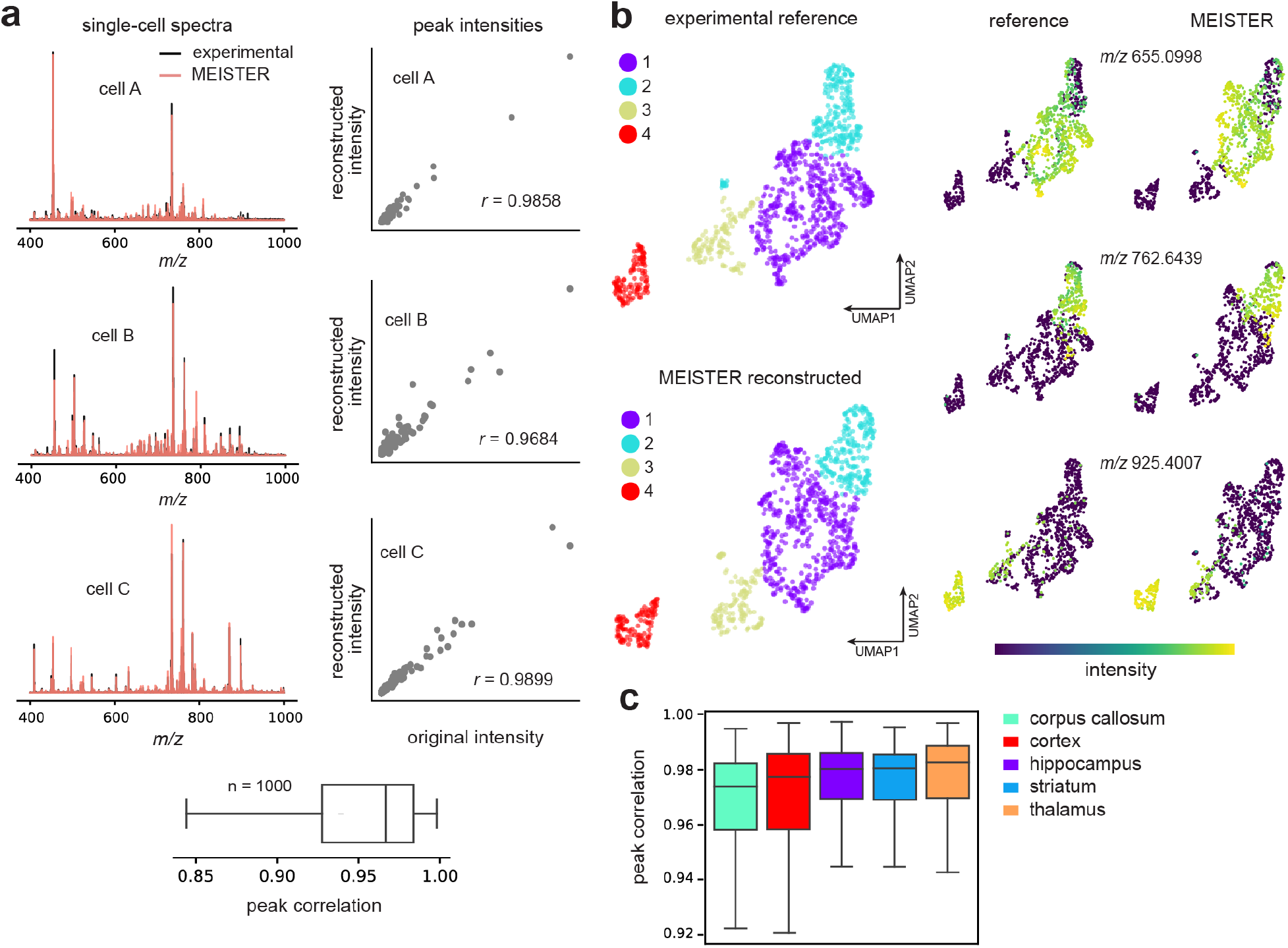
**a**, Experimental high-resolution single-cell mass spectra versus MEISTER reconstructed mass spectra. High peak correlation scores were obtained on 1000 validation cells (bottom box plot). **b**, Downstream analysis of the reference (full transients) and reconstructed data shows nearly identical clustering patterns through k-means (k=4; 1-4 denote cluster numbers and each cluster of cells are coded with a different color) and ion distributions at the single-cell level. **c,** Peak correlation scores for cells sampled from five different brain regions.

**Extended Data Fig. 4.**
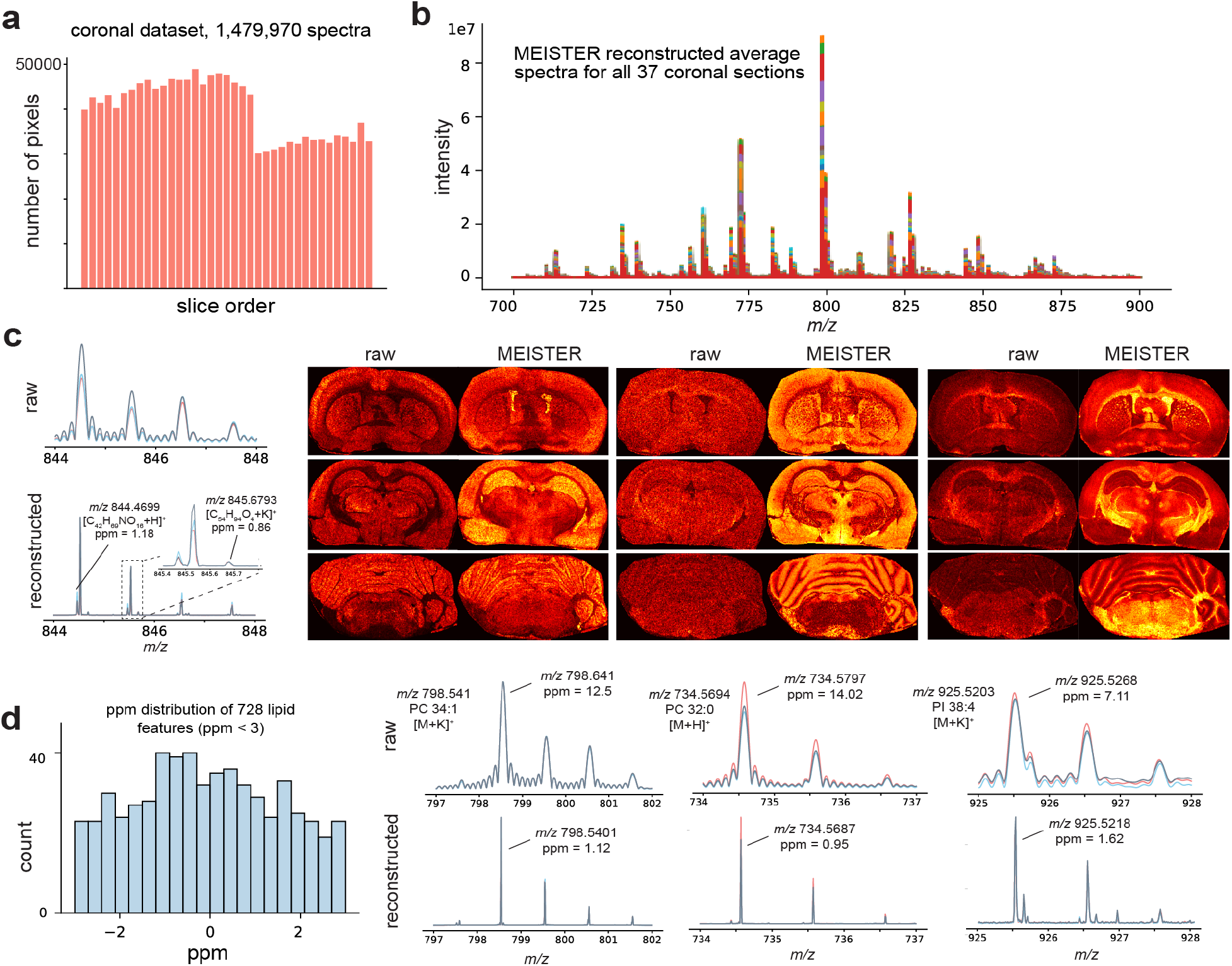
**a**, Number of pixels versus the slice order for the 3D rat coronal data set. **b**, Average mass spectra for 37 coronal sections obtained from MEISTER reconstruction. **c,** Comparison of the raw (reduced) and reconstructed mass spectra (left) in a small *m/z* window, and representative ion images (right). **d**. Distribution of ppm mass errors of 728 matched lipid features (left) and comparisons of mass spectra and mass resolution for several common brain lipids (right).

**Extended Data Fig. 5.**
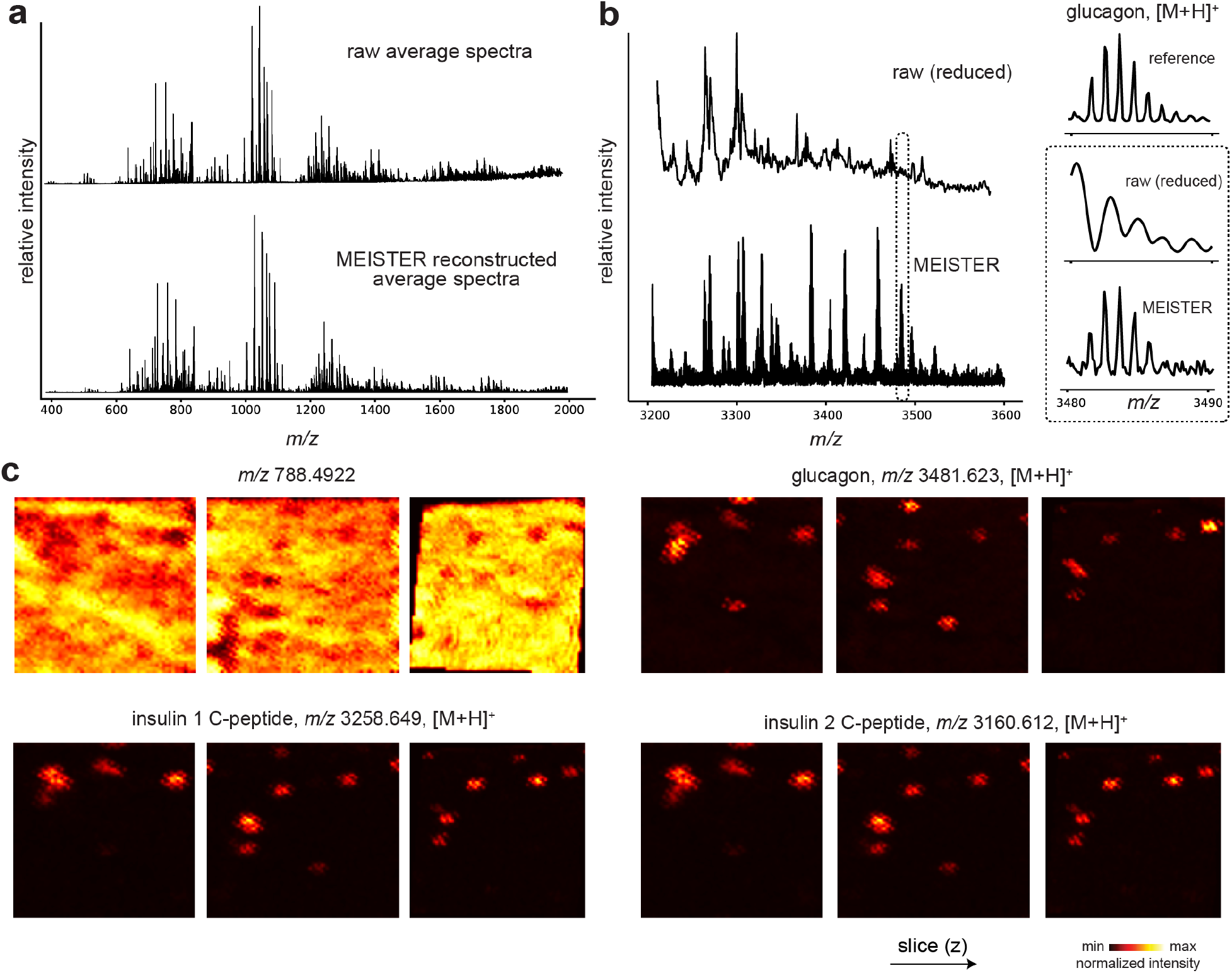
**a,b**, Averaged mass spectra of rat pancreas tissue sections from the reduced data (top) and our reconstruction (bottom) for *m/z* range of 400 to 2000 (a; lipids) and 3200 to 3600 (b; peptides). Inlet displays a zoomed-in *m/z* window with signals of protonated glucagon for high-resolution reference (top), reduced (middle) and reconstructed (bottom) data. High-fidelity reconstruction by the proposed method w.r.t. the reference can be observed. **c**, Ion images of different sections obtained from deep learning reconstructed tissue MSI data for *m/z* 788.4922, glucagon, insulin 1 C-peptide, and insulin 2 C-peptide.

**Extended Data Fig. 6.**
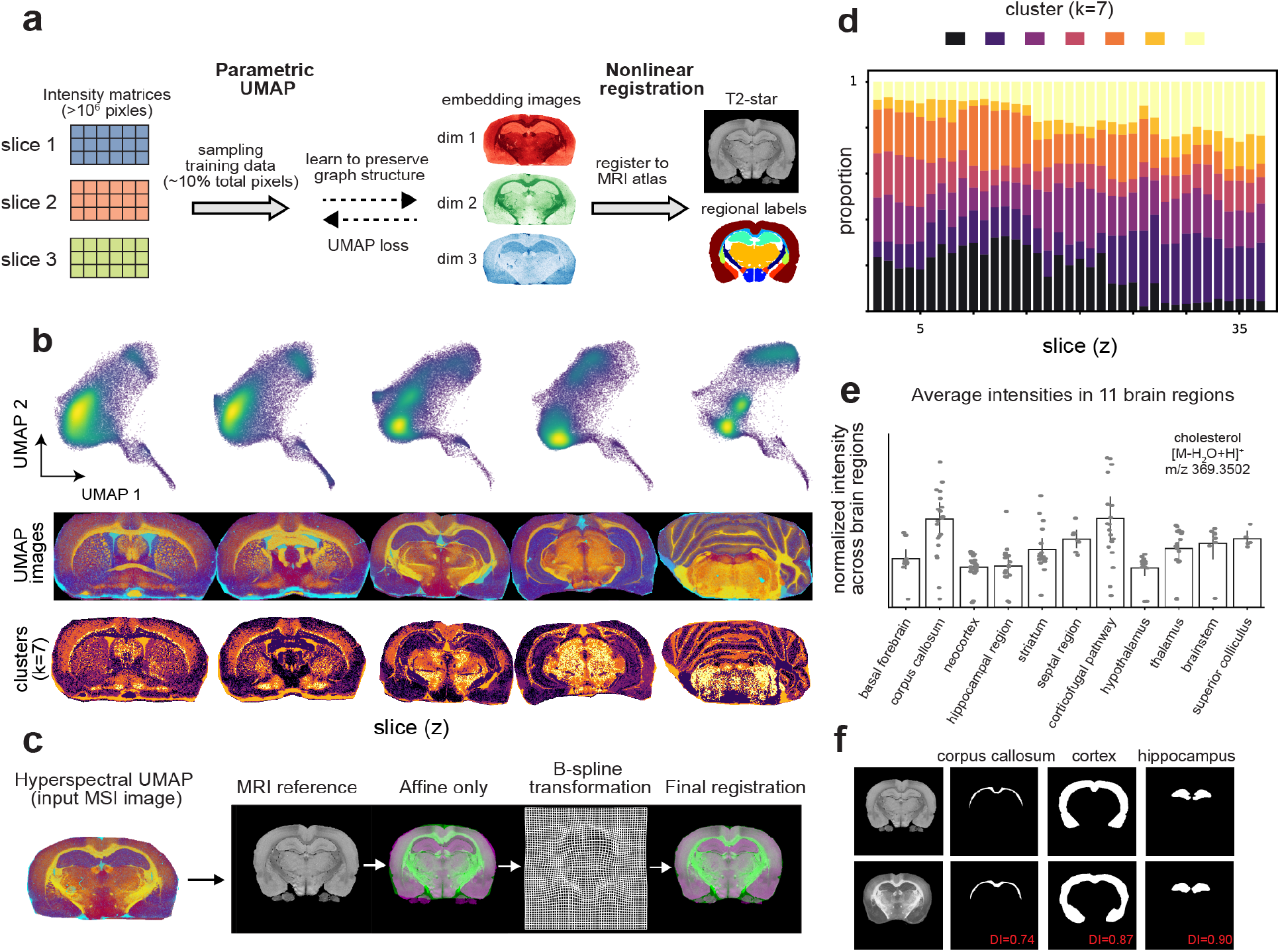
High-resolution 3D MSI of the brain with volumetric reconstruction. **a**, The proposed workflow leveraging pixel-wise parametric UMAP to efficiently align MSI data from serial tissue sections to a 3D MRI volume. **b,** Across serial sections in z-axis, parametric UMAP embeddings (top, colored by point density) formed consistent and structurally informative feature images (middle) and k-means clusters (bottom), **c,** Nonlinear image registration aligning the hyperspectral UMAP image to the target MR image via sequential affine and B-spline registration. The combination produced excellent alignment and enabled coherent volumetric reconstruction of MSI sections. **d**, The cluster proportions for 7 clusters varying with the slice order. **e**, Differential signal intensity distributions for lipids across 11 brain anatomical structures identified using the ROI labels from a rat MRI atlas, i.e., cholesterol shown here with distinct regional differences. **f**, Anatomical masks from the atlas (top) and manual annotations of MSI post registration (bottom). Dice Indices are shown in red indicating good alignment.

**Extended Data Fig. 7.**
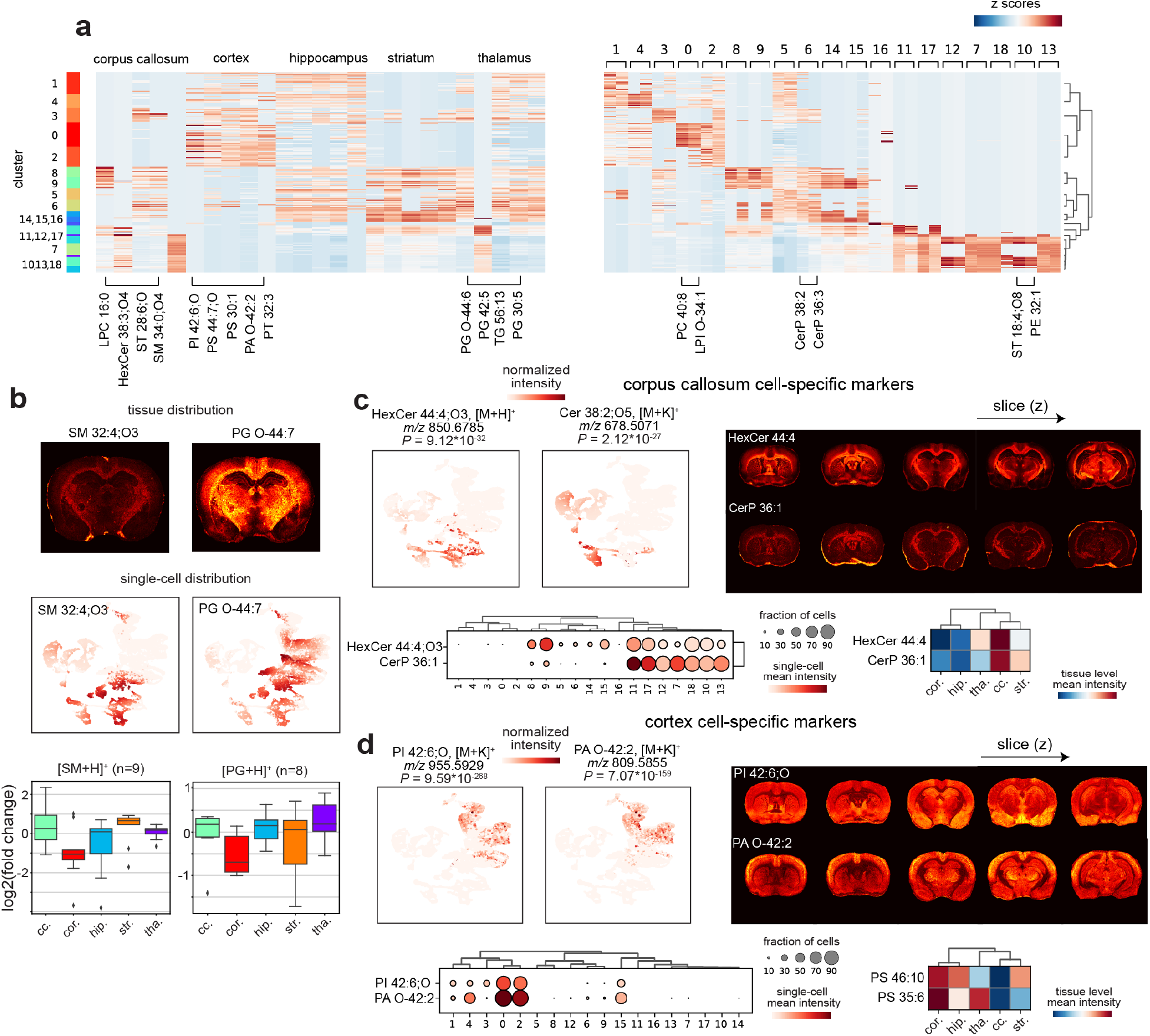
**a**, Top five single-cell lipid features identified to be brain-region specific (left), and top two lipid features identified to be cluster-specific (right). Lipid features were selected based on p-values and log2 of fold change obtained by differential analysis. Rows and columns correspond to cells organized by the clusters and lipid features organized by regions respectively. Region and cluster specific lipids can be identified. For instance, PI 42:6;O is significantly elevated (adj. p-value=9.6*10^-268^) in cells from cortex. Inspecting the PI 42:6;O column in the heatmap, we can observe that most cells in cluster 0, 2, 3, 4 contain this particular lipid. **b**, Tissue and single-cell distributions of lipid markers identified by region-specific lipid analysis, demonstrated by data of two representative lipids, SM (32:4);O3 and PG O-(44:7). Bottom: regional distributions of SM and PG signal intensities quantified by log2 of fold change at the single-cell level. **c, d,** Highly-specific lipid markers (significance indicated by p-values) were identified for different brain regions, showing agreement between single-cell and tissue imaging data for **c,** corpus callosum and **d,** cortex regions. For **c, d,** top left: single-cell UMAP, top right: corresponding ion images, bottom left: relations between clusters, mean signal intensity for single cells in cluster, and size of cell fraction per cluster, bottom right: relation between mean of signal intensities, brain locations of collected signals.

**Extended Data Fig. 8.**
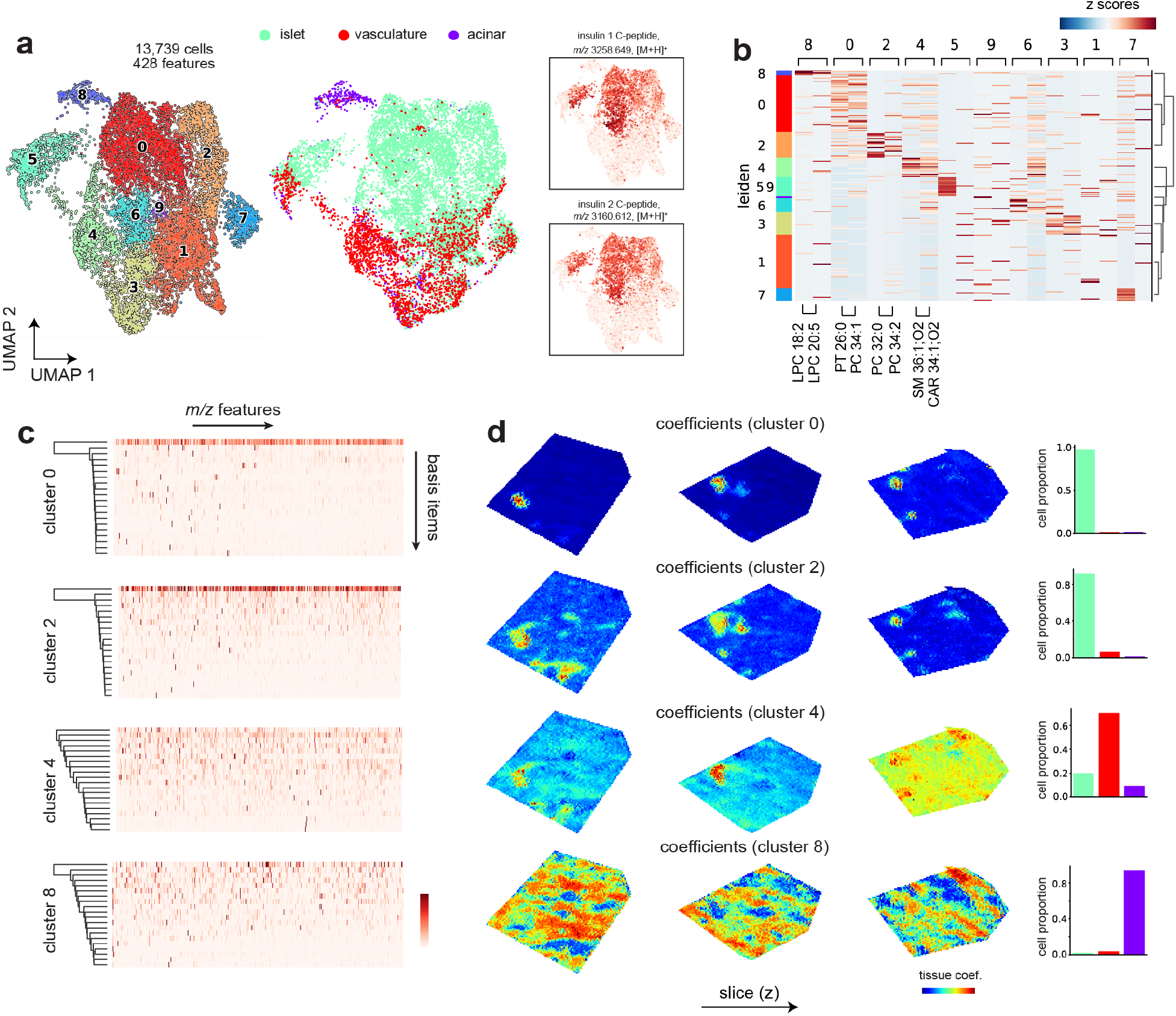
**a**, A total of 13,739 cells from rat pancreas with 428 cross-annotated features (tissue and single cells) are subjected to UMAP and Leiden clustering analysis. 10 cell clusters were identified (left), which can also be mapped to three major pancreatic cell types (right). Inlet shows the distributions of insulin 1 and 2 C-peptides within single cells. **b**, Top two features identified to be cluster-specific across all clusters. **c**, Cell-cluster-specific dictionaries extracted from representative cluster 0, 2, 4, and 8. **d**, Estimated spatial contributions of individual cell clusters across pancreas tissue. Each row shows results of mapping the contributions of one cluster to individual pixels, revealing distinct spatial organizations of islet, vasculature, and acinar cells.

## SUPPLEMENTARY INFORMATION

**Supplementary Figure 1.**
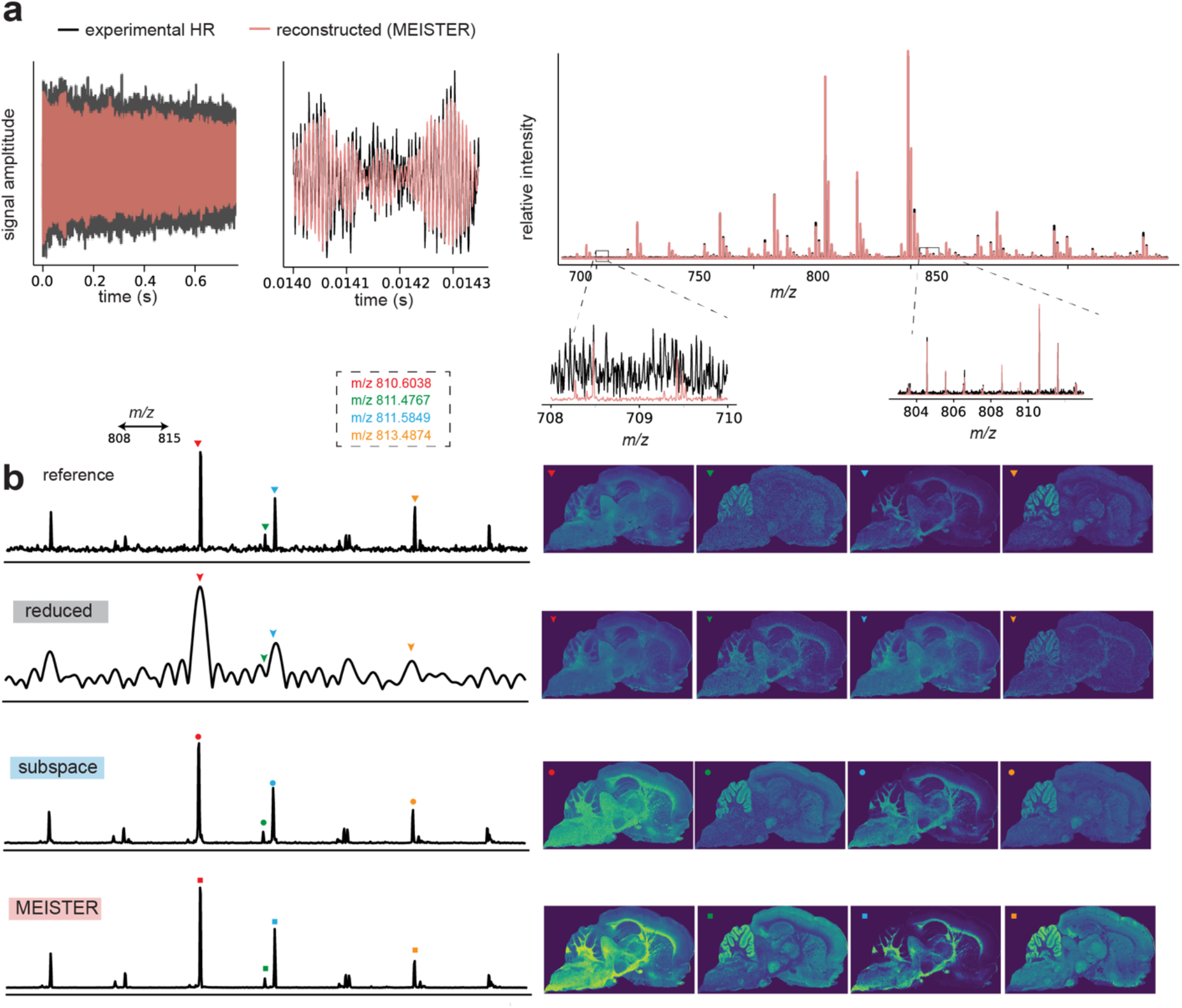
**a**, Experimental and reconstructed transients (1M temporal points for 0.731 s duration) and the transformed mass spectra. **b**, From top to bottom rows: reference, reduced, subspace and MEISTER reconstructed mass spectra (left column), and ion images formed from signals indicated by the colored markers (right column). The data were from a validation data set of a sagittal section adjacent to the training section.

**Supplementary Figure 2.**
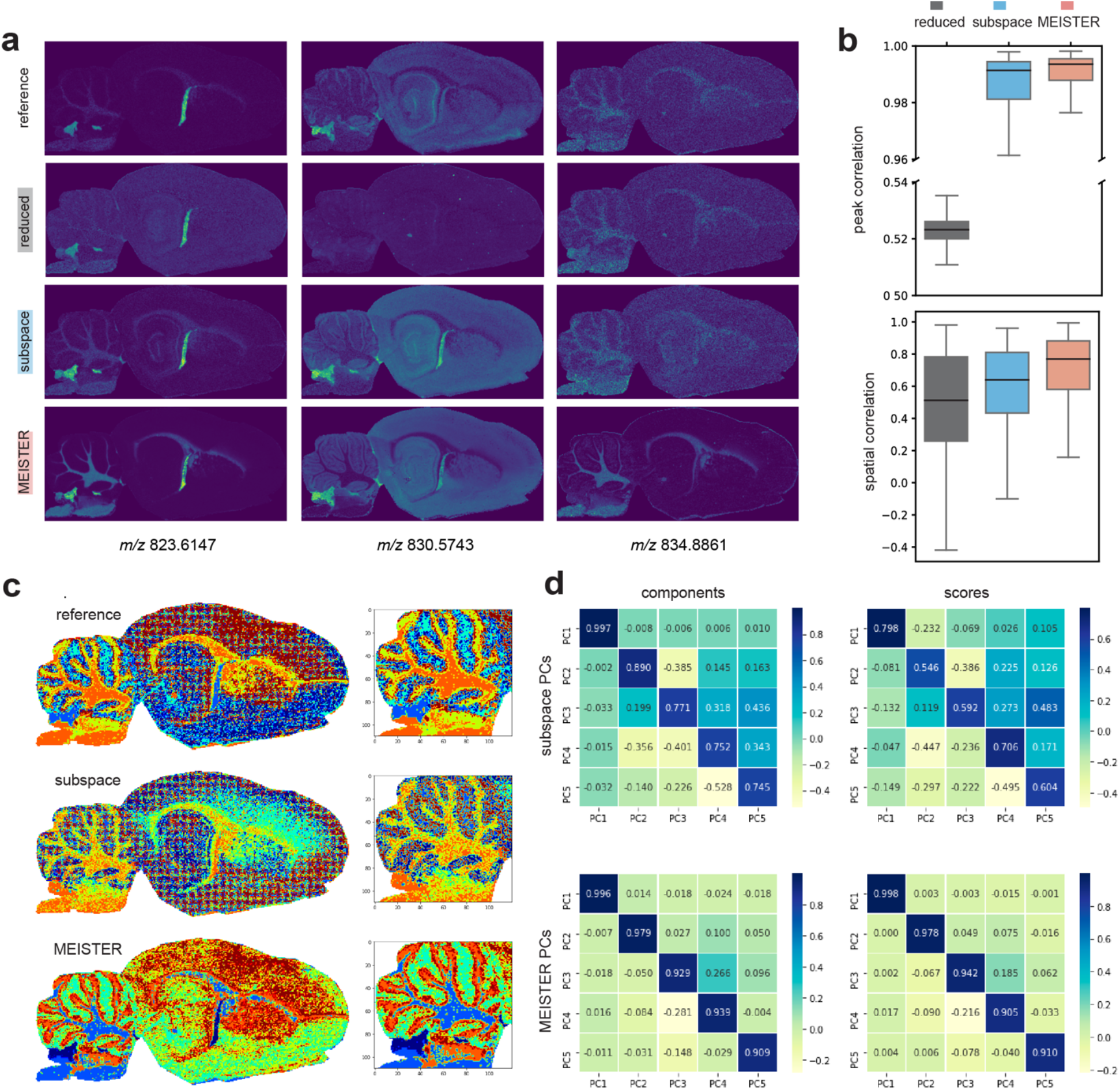
**a**, From top to bottom rows: reference, reduced, subspace, and MEISTER reconstructed ion images from a validation data set acquired on a sagittal section 2 mm away from the training section. **b**, Peak and spatial correlation scores compared to the reference. **c**, K-means clustering assignments for the reduced and reconstructed data by the subspace approach and MEISTER. **d**, Pearson correlation coefficients between top-5 PCs extracted from the experimental reference versus from the data reconstructed by subspace (top) and MEISTER (bottom).

**Supplementary Figure 3.**
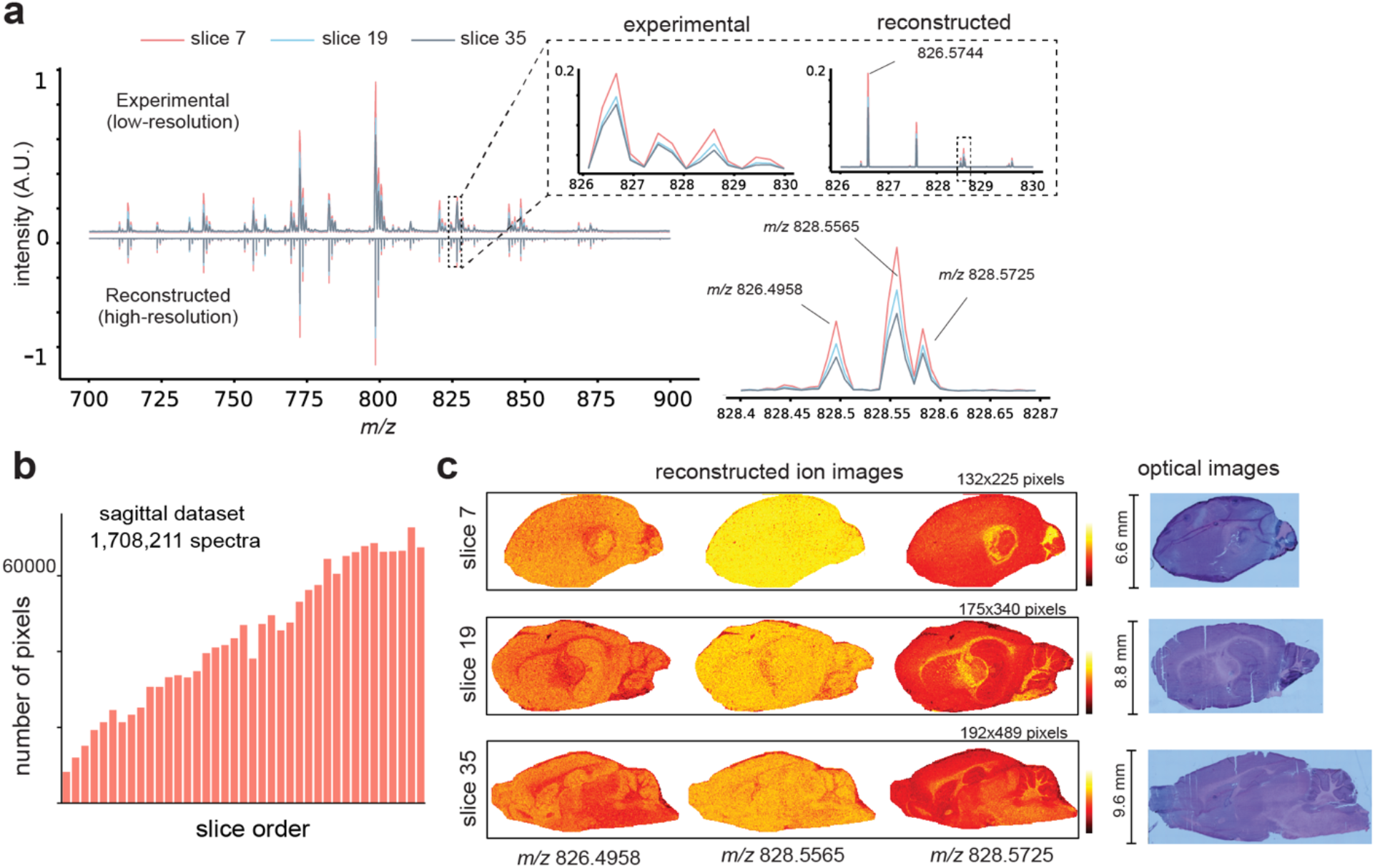
Reconstruction of the 3D sagittal data set showing a, Average mass spectral from 3 sagittal sections obtained from MEISTER reconstruction. **b,** Number of pixels versus the slice order for the 3D rat sagittal data set. **c,** Ion images reconstructed by MEISTER (left) and the optical images from the same tissue sections.

**Supplementary Figure 4.**
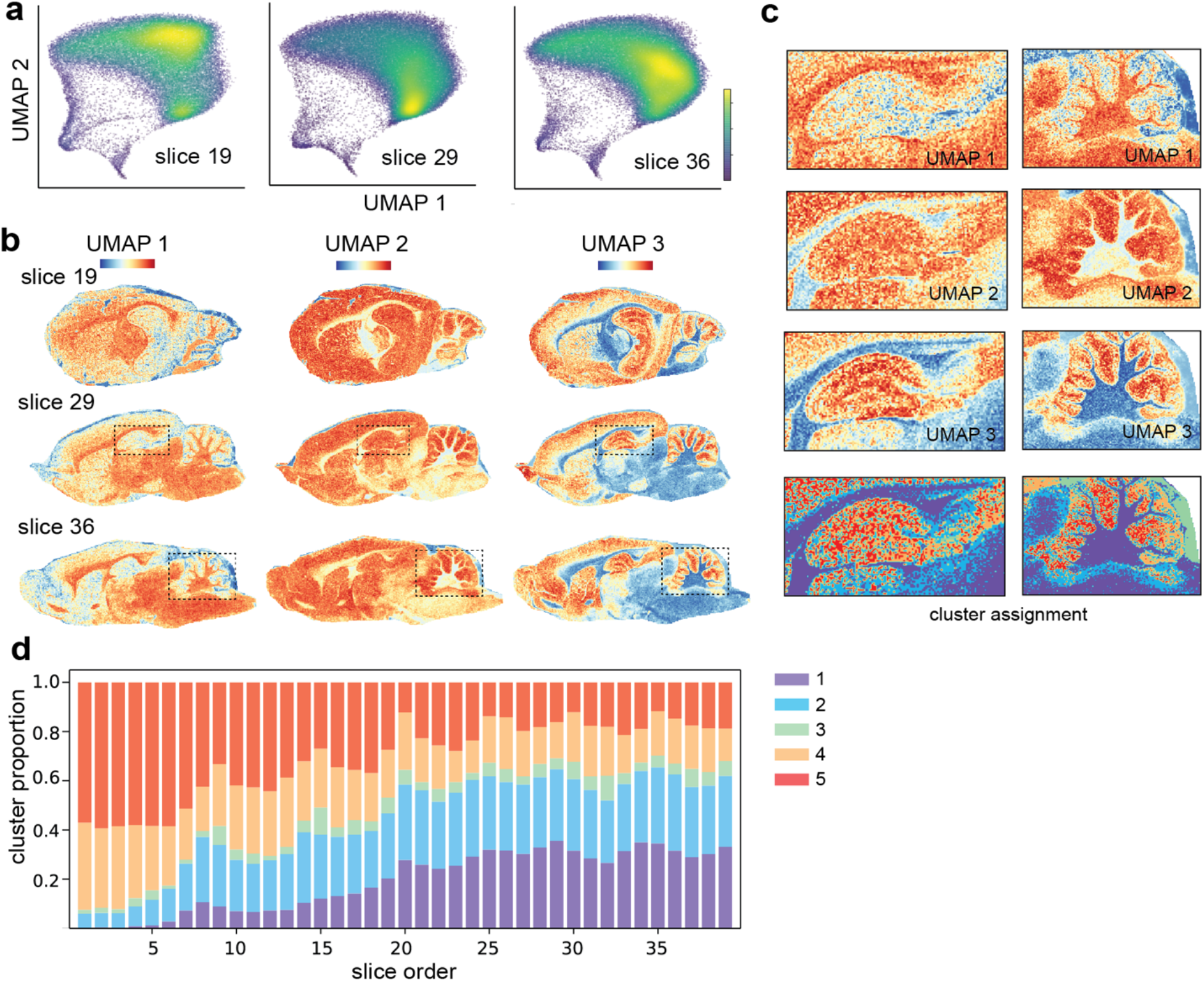
**a**, Parametric UMAP embeddings of the 3D sagittal data set, data shown on 3 representative sections. **b,** Feature images formed by individual UMAP dimensions. **c,** Zoom-in views of the feature images and the k-means clustering assignments, with **d** showing the cluster proportions for 5 clusters varying with the slice order.

**Supplementary Figure 5.**
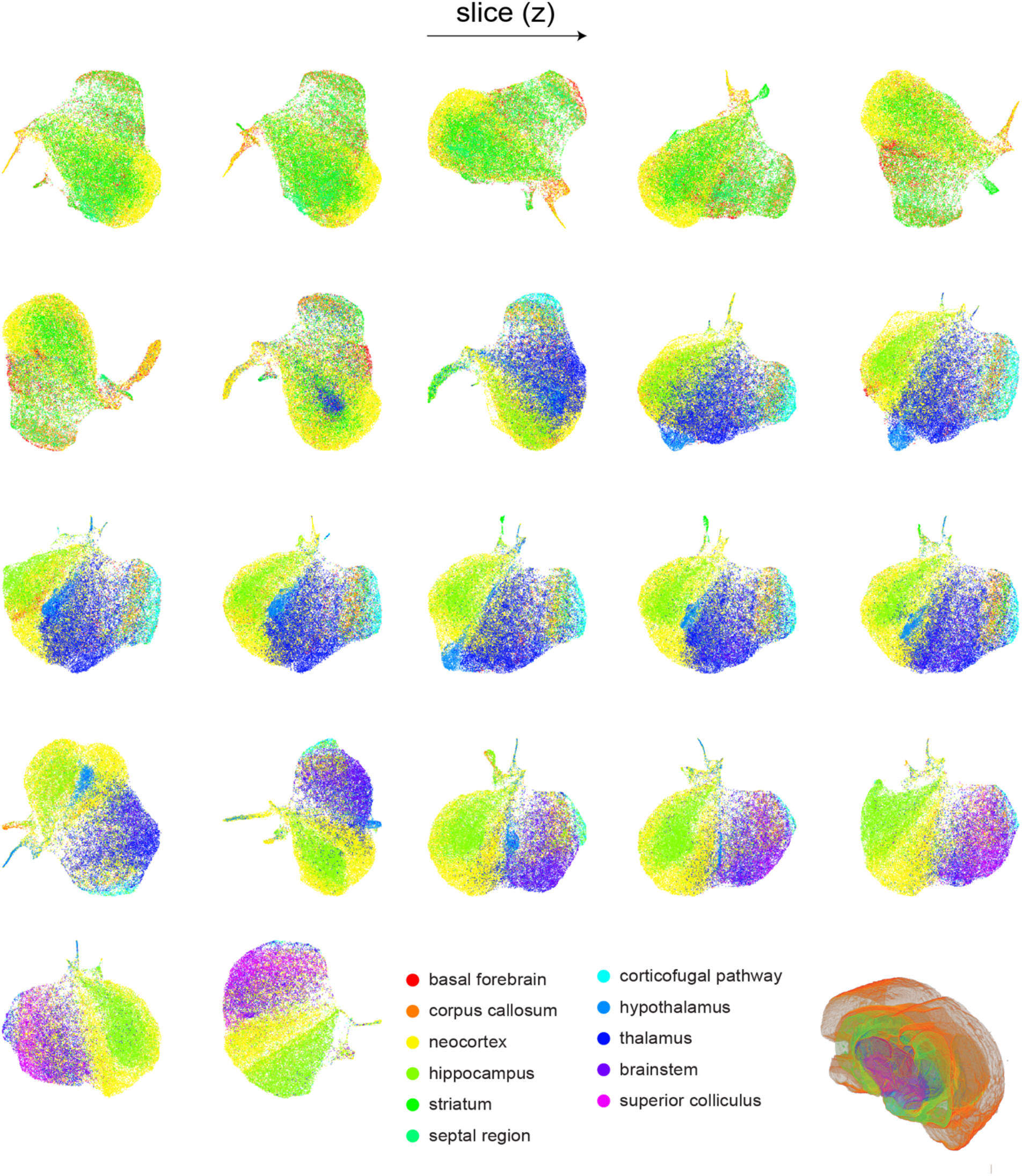
UMAP embeddings of representative 2D tissue sections covering all 11 brain regions (colors indicated at the bottom).

**Supplementary Figure 6.**
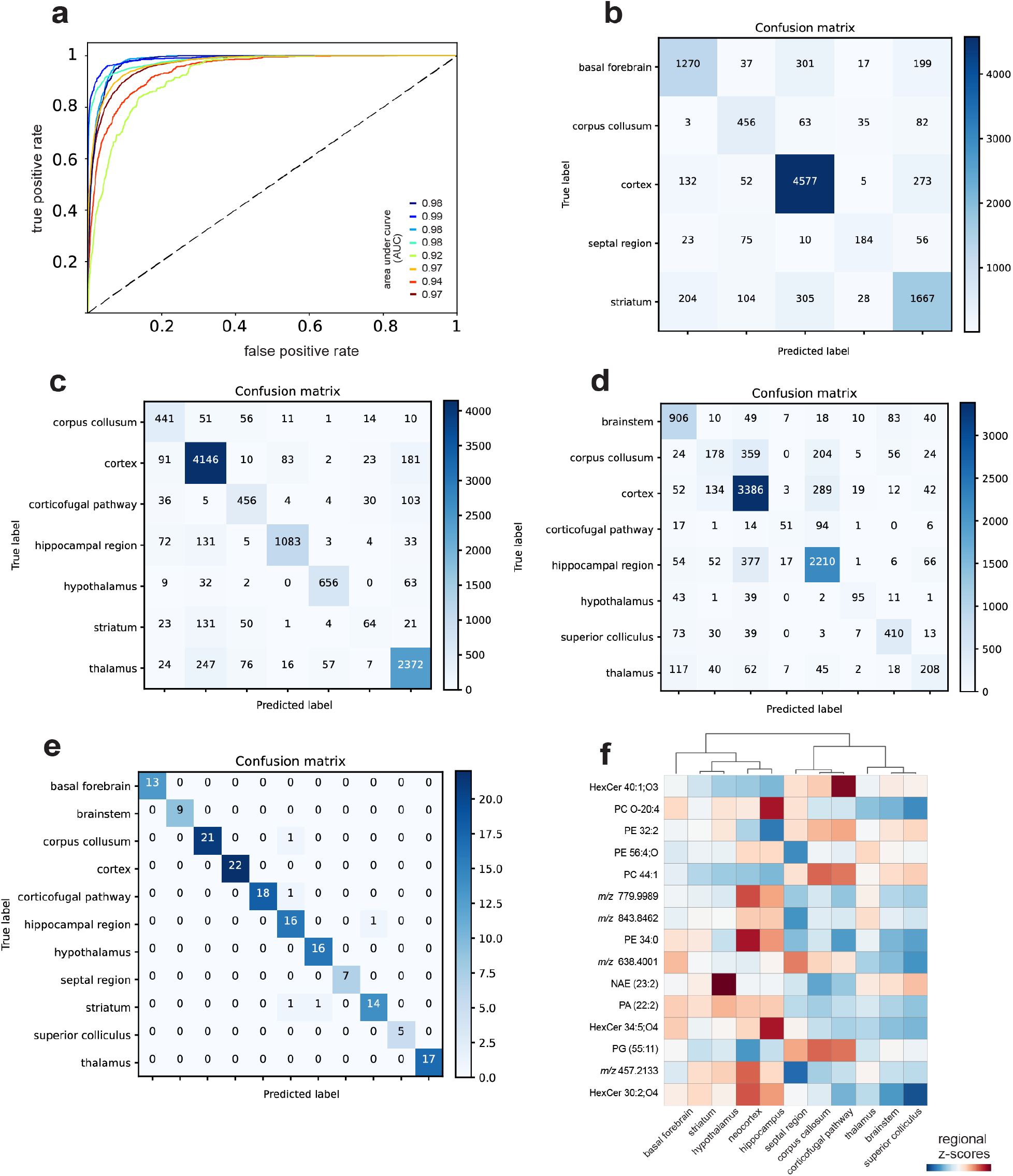
Machine learning models trained on 3D MSI pixels to predict brain regions. **a,** The AUC curves of a model trained on a tissue section containing 8 brain regions. **b-d,** Confusion matrices (validated on 20% of pixels) of the models for three tissue sections demonstrated in Figure 4 (top left, top right, and bottom left), and **e,** the confusion matrix of the model trained on the average mass spectra (bottom right). **f**, Mean intensity profiles of the top selected lipid features for different brain structures.

**Supplementary Figure 7.**
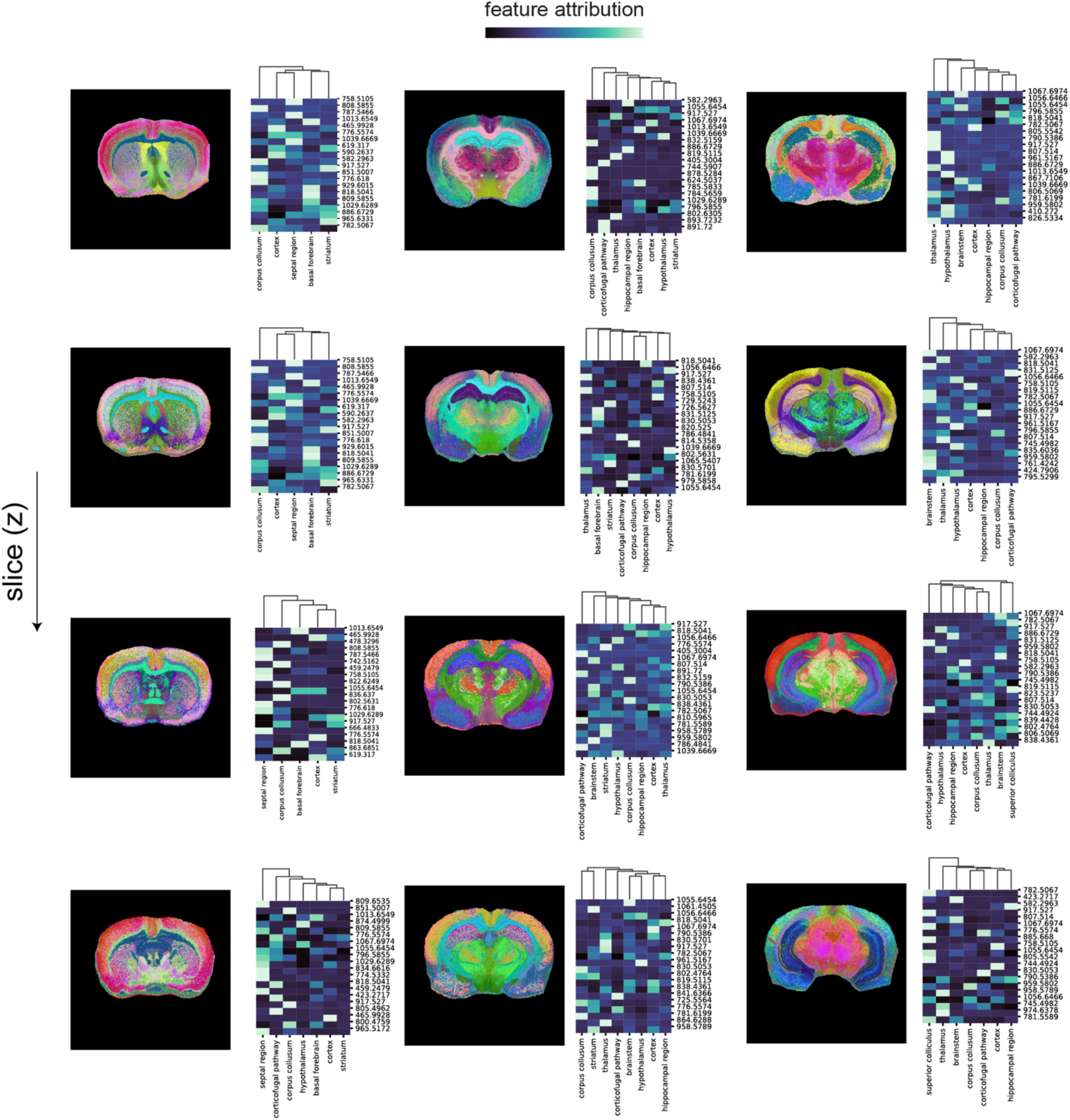
Machine learning models trained on individual MSI pixels to predict brain regions and using interpretable machine learning based on SHAP values for top contributing lipid features toward model predictions. For each tissue section, image representations of the UMAP on SHAP values (left) and average SHAP values of top-20 lipid features (right) are shown.

**Supplementary Figure 8.**
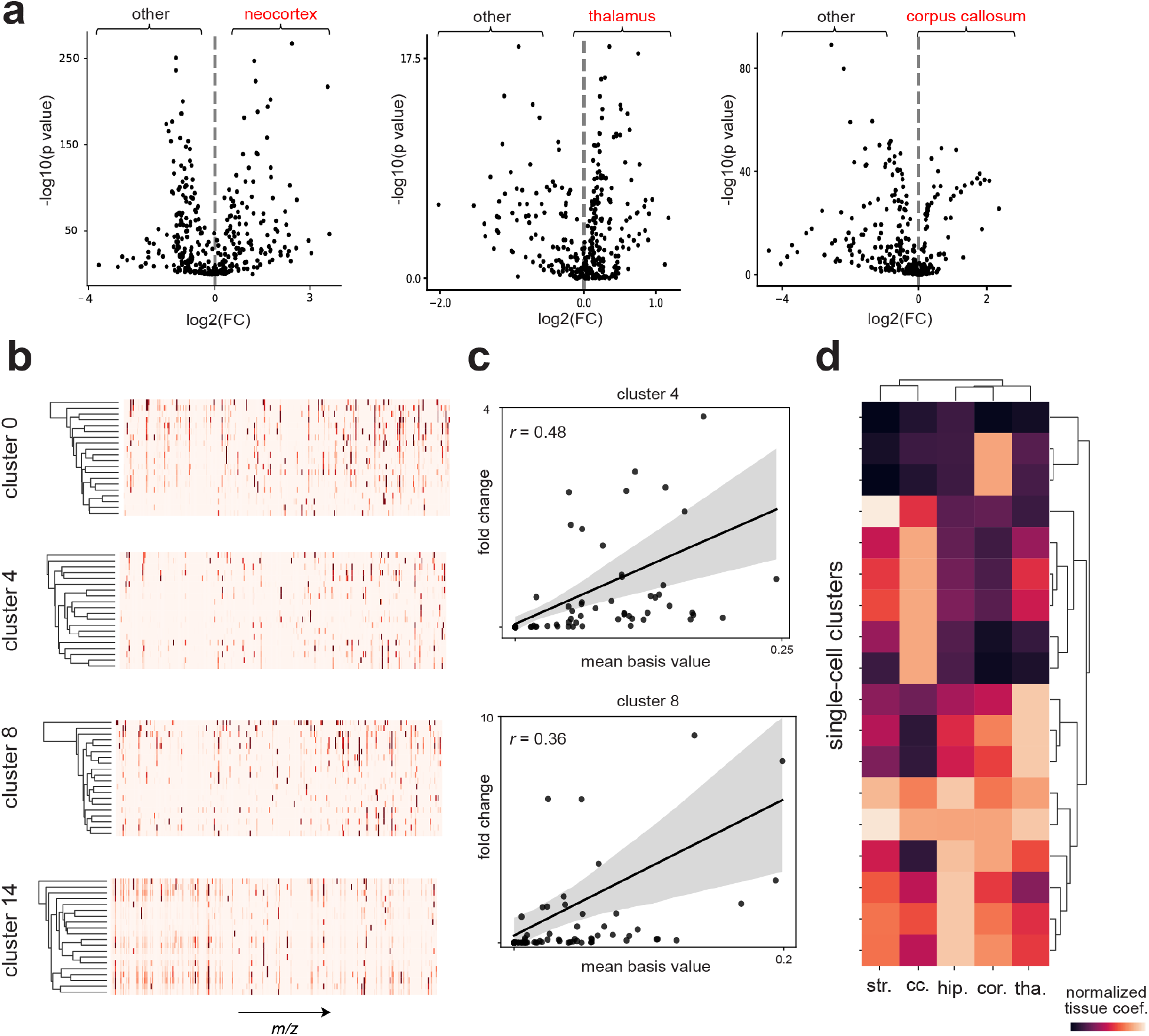
**a**, Volcano plots obtained from differential analysis of regional-specific lipids for 13,566 single cells. **b**, Additional representative cell-specific chemical dictionaries extracted from clusters. **c**, basis values averaged on 20 dictionary items versus fold change for lipid features in cluster 4 and 8. **d**, Analysis of contributions of cell-specific biochemical signatures to brain regions, by summarizing the estimated pixel-wise model weights for each single-cell clusters. Distinct cell compositions can be observed for different regions.

**Supplementary Figure 9.**
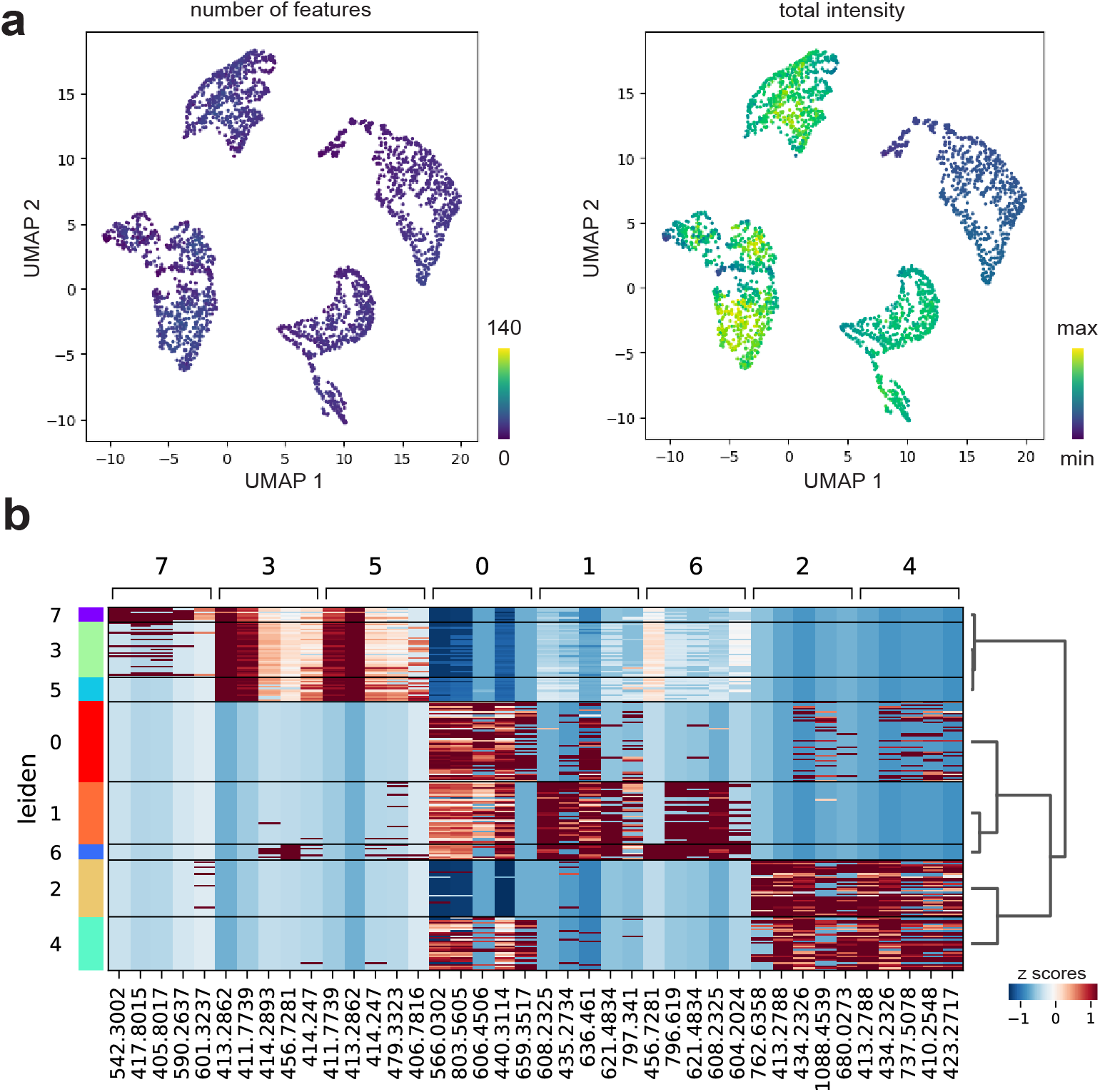
**a**, UMAP of 2,692 hippocampal cells overlaid with number of lipids per cell (left) and total intensity per cell (right). **b**, Heatmap showing the normalized scores (z scores) for the top-5 differential *m/z* features of 8 hippocampal cell clusters.

**Supplementary Figure 10.**
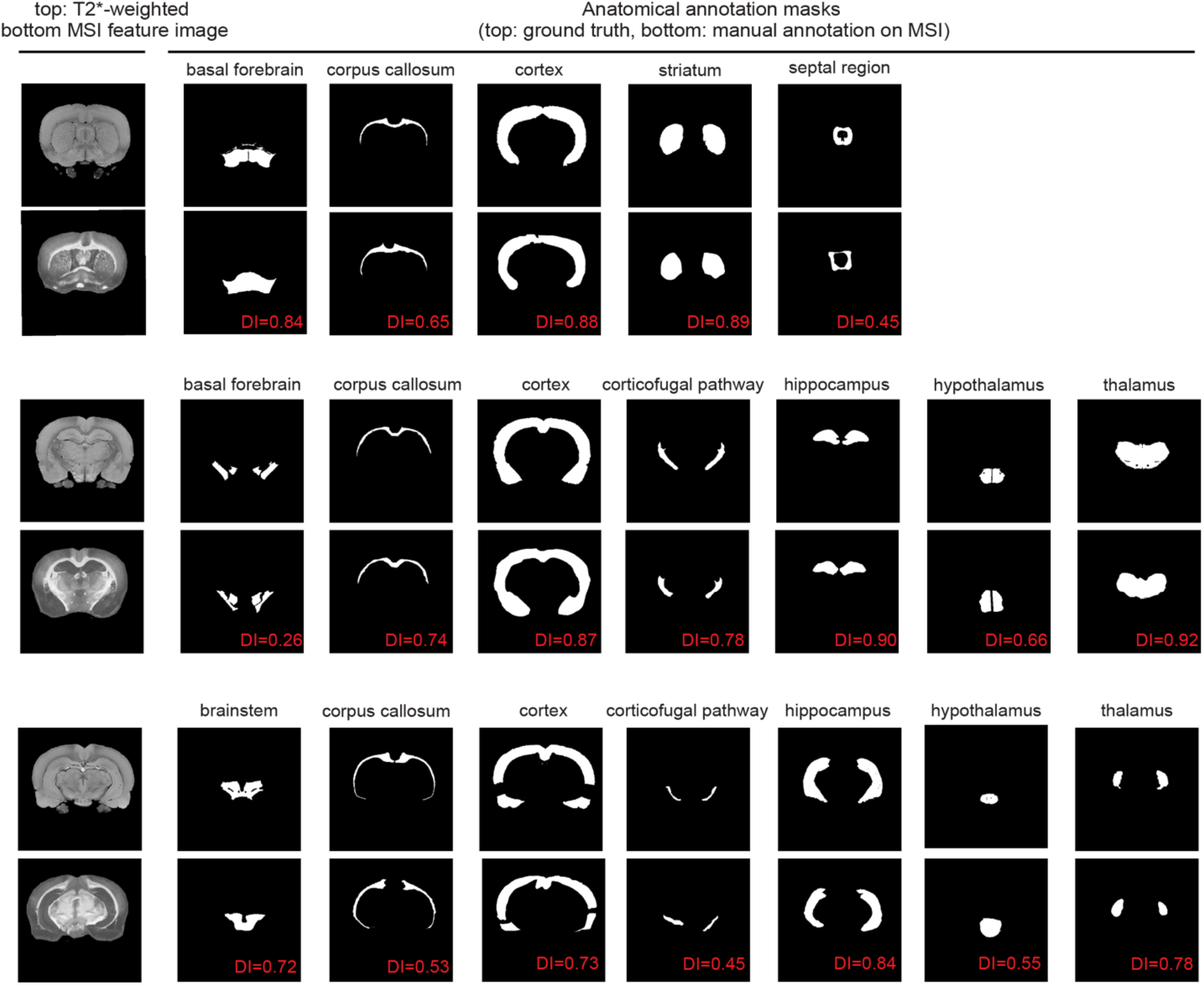
Group truth (atlas) and manual annotated (registered MSI) masks for different brain regions. Dice Indices computed for each mask pairs were indicated in red.

